# An electrical brain network encodes anxiety in health and depressive states

**DOI:** 10.1101/2024.06.26.600900

**Authors:** Dalton N. Hughes, Michael Hunter Klein, Kathryn Katsue Walder-Christensen, Gwenaëlle E. Thomas, Yael Grossman, Diana Waters, Anna Matthews, William E. Carson, Yassine Filali, Mariya Kaminsky, Alexandra Fink, Neil M. Gallagher, Masiel Perez-Balaguer, Colleen A. McClung, Jean Mary Zarate, Rainbo C. Hultman, Stephen D. Mague, David E. Carlson, Kafui Dzirasa

## Abstract

In rodents, anxiety is characterized by heightened vigilance during low-threat and uncertain situations. Though activity in the frontal cortex and limbic system is fundamental to supporting this internal state, the underlying network architecture that integrates activity across brain regions to encode anxiety across animals and paradigms remains unclear. Here, we utilize parallel electrical recordings in freely behaving mice, multiple translational paradigms known to induce anxiety, and machine learning to discover a multi-region network that encodes the anxious brain state. The network is composed of circuits widely implicated in anxiety behavior, it generalizes across many behavioral contexts that induce anxiety, and it fails to encode multiple behavioral contexts that do not. Strikingly, the activity of this network is also principally altered in two mouse models of depression. Thus, we establish a network-level process whereby the brain encodes anxiety in health and disease.

---

Anxiety is a mental state marked by heightened pressure, concern, or apprehension related to uncertain future circumstances ^1^. The anxious state can be adaptive to increase the rate of survival, or it can become overly generalized and persistent in a manner that yields behavioral pathology that can lead to anxiety disorders. These disorders constitute the largest group of mental disorders in Western and high-income societies, with nearly 34% of U.S. adults directly impacted in their lifetime ^1,2^. Strikingly, the prevalence of symptoms of anxiety disorders or other mental health disorders increased during the height of the COVID-19 pandemic ^3^. As such, it is imperative to discover the biological basis of the anxious brain state and to delineate how the brain encodes anxiety in the disordered state.

Non-invasive human imaging studies have demonstrated altered activity in multiple cortical and limbic brain regions, including the amygdala, prefrontal cortex, and hippocampus during heightened anxiety, as well as synchronized activity between these regions and others at the millisecond-to-second timescales ^4-7^. Human intracranial recordings have also linked higher trait anxiety to altered coherence in networks with some of these regions ^8^, pointing towards the involvement of integrated multi-regional circuits in mediating the anxiety state.

A myriad of rodent studies have implicated homologous regions in mediating anxiety: the amygdala (Amy), ventral hippocampus (VHip), and subregions of the medial prefrontal cortex (mPFC). Pharmacological lesions and optogenetic inactivation studies have implicated the necessity of these brain regions for anxious behaviors ^9,10^. Furthermore, precise circuit-level studies in rodent models have further delineated the role of these brain regions and their integrated circuits ^11^. For example, millisecond-level coherence is observed between the mPFC, Amy and/or VHip during key aspects of anxiety-related behaviors ^12-17^, and optogenetic interrogation of projections involving these regions modulate anxiety-related behaviors ^18-23^. Yet it remains to be clarified how these circuits reliably integrate across timescales (i.e., network-level stability) to selectively encode anxiety across animals and behavioral contexts (i.e., generalization) in healthy animals and in disease states.

Because local field potentials (LFPs) capture generalized patterns of neural activity that can be consistently sampled across subjects ^24^, we previously developed a machine learning technique called discriminative Cross-Spectral Factor Analysis-Nonnegative Matrix Factorization (dCSFA-NMF) to discover behaviorally relevant ensembles of LFP activity that synchronize at both the milliseconds and seconds timescale [i.e., electrical functional connectome (’electome’) – networks]. An electome network can be composed by LFP oscillatory power from each brain area, millisecond-resolution coherence between oscillations from pairs of brain regions, and/or directionality (an indication of information transfer between pairs of brain regions assessed using Granger causality testing), ranging from 1–56Hz. Moreover, dCSFA-NMF was designed to discover electome networks that encode behaviorally relevant internal states both within and across mice ^25-27^.

Here, we operationalized anxiety as an internal state characterized by increased behavioral inhibition in response to perceived threat, which can be assessed through measurable, test-dependent outcomes. These can include (but are not limited to) reduced exploration of exposed spaces, heightened physiological responses, hyperarousal, and vigilance patterns that facilitate the association of an ongoing threat with the context in which it is experienced. We then used dCSFA-NMF to discover a distinct electome network that selectively encodes anxiety induced by multiple contexts in healthy animals. The network was also activated in mouse models of stress disorders in a context that should otherwise not promote anxiety.

## RESULTS

### Distributed electome networks encode a convergent anxious internal state

Forty-one male mice were implanted with multiwire electrodes to concurrently target mPFC [cingulate (Cg), prelimbic (PrL) and infralimbic (IL) cortex], Amy, VHip, nucleus accumbens (NAc), medial dorsal thalamus (MD), and ventral tegmental area (VTA; Fig. 1A, left). Following their recovery, we employed a two-stage approach to discover how distributed neural activity encodes an internal state for anxiety. First, we utilized a translational anxiogenic protocol based on treatment with the antidepressant fluoxetine (FLX). Acute administration of this class of agents is known to induce anxiety in humans, and it has been shown to increase behavioral inhibition in response to perceived threat in mice ^28^ relative to a saline control condition (see also Supplemental Figure S1).

**Figure 1:**
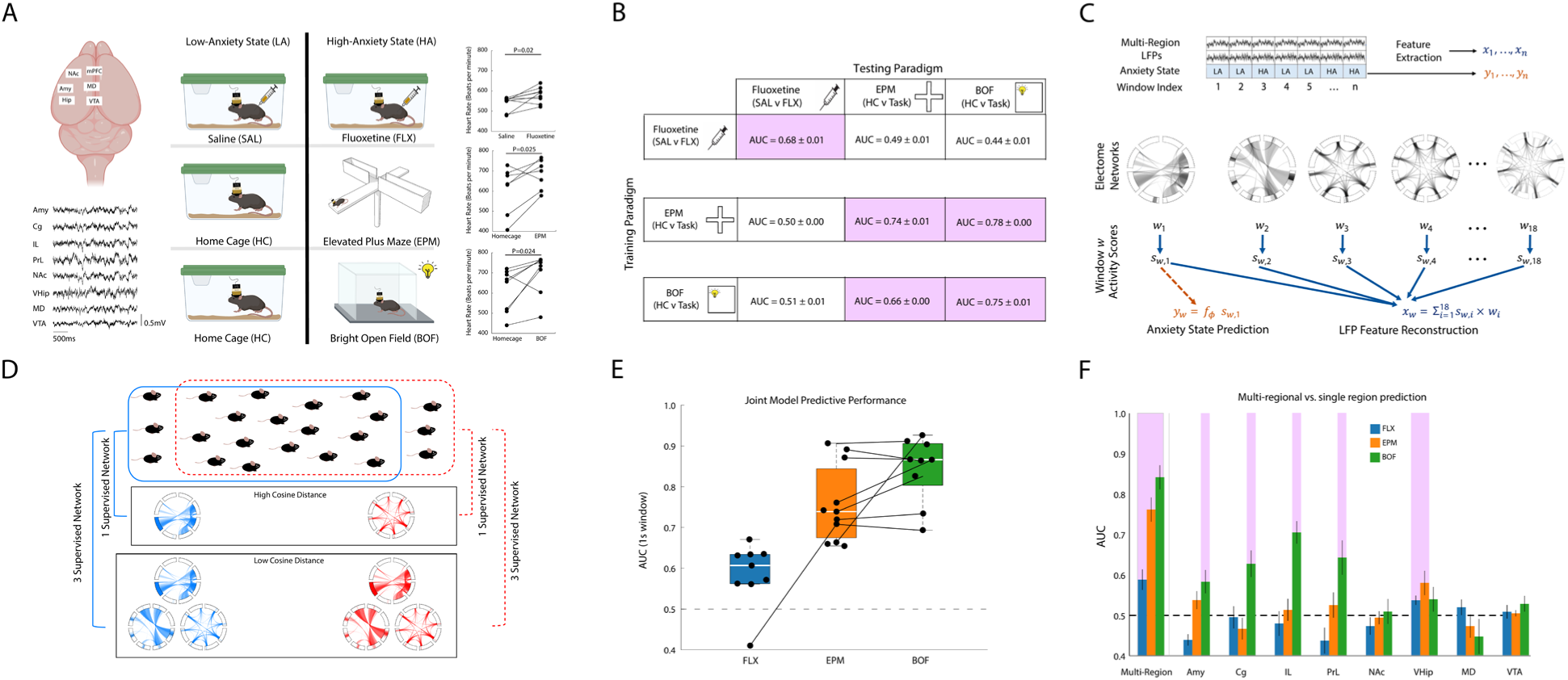
Distributed electome networks encode anxiety states across multiple anxiety-related paradigms. **A)** Local field potential oscillations recorded concurrently from 8 brain regions (left), as mice were subjected to three distinct paradigms used to induce low- (LA) and high-anxiety (HA) states respectively (middle column): acute injections of saline vs. fluoxetine (top), home cage vs. elevated plus maze (middle row), and home cage vs. bright open field (bottom). Heart rate measures for each assay are shown to the right, and comparisons between the LA and HA states were performed using a one-tailed t-test (P<0.05 for all comparisons, N=7-9 male mice per assay). **B)** dCSFA-NMF results when the network model was used to discover an electome network for each anxiety paradigm. Electome networks learned for the three anxiety paradigms were applied to new mice subjected to the three paradigms (N= 13, 26, and 19 training mice for FLX, EPM and BOF, respectively, and N= 6, 11, 9 holdout testing mice for FLX, EPM and BOF, respectively). Nine generalization tests for each of the three learned networks were run in new mice subjected to the three different anxiety paradigms. AUC value of 0.5 corresponds to models with no predictive utility. **C)** Schematic of electome-based decoding of anxiety from multi-region LFP activity. Multi-region LFP recordings are segmented into discrete time windows that are labeled as low- or high-anxiety states and converted into feature vectors describing oscillatory power, coherence, and directionality (top). These features are decomposed into a set of electome networks (middle), producing window-specific network activity scores (bottom). A supervised model maps a subset of the network activities to an anxiety state for each window (bottom left), while the learned network weights allow reconstruction of LFP features from network activity (bottom right), providing interpretability of the latent network representations. **D)** Network consistency was evaluated by training the dCSFA-NMF model multiple times, where the mice used for training and validation were shuffled. A cosine distance metric quantified the consistency of the supervised networks across runs, where a lower cosine distance reflected greater network consistency. **E)** Box and whisker plots show generalization tests in which the networks learned from the multi-assay dCSFA-NMF model were applied to new mice (same holdout testing mice as Fig. 1B) subjected to the three different anxiety paradigms. Horizontal dashed line at AUC = 0.5 corresponds to models with no predictive utility. For the box and whisker plot: central mark is the median, box edges are the 25th and 75th percentiles, whiskers extend to extreme data points that are not outliers, and outliers are plotted individually as "+." Solid lines indicate the data points from the same mouse. **F)** Predictive utility of multi-region, multi-assay dCSFA-NMF network model (same as Fig. 1E) vs. models solely based on activity from single brain regions. Models that showed significant assay encoding are highlighted in pink (single-sample t-test against a null AUC distribution at α = 0.05, shown as mean±s.e.m). Note that only the network model encoded all three assays.

Second, we tested whether the electome networks learned for this paradigm also encoded the internal state induced by two paradigms characterized by increased behavioral inhibition with regards to exploration of exposed spaces: the elevated plus maze (EPM) and the bright open field (BOF). The amount of time a mouse spends within exposed regions of these assays (open arms in the EPM and center of the BOF) is widely used to infer anxious internal states. As mice with the highest anxiety-like behavior may never enter the exposed regions of these assays, we modeled high anxiety as the internal state that was causally induced by the three paradigms (Fig. 1A, middle). All three paradigms heightened physiological responses in mice in the high-anxiety context, as assessed via heart rate (T_6_=2.6, T_7_=2.4, T_8_=2.9 and P=0.025, P=0.024, P=0.01 for the EPM, BOF, and FLX assays, respectively, one-tailed t-tests; N=7-9 male mice per assay, see Fig. 1A, right), as expected based on our operational definition of anxiety. We trained our network model at one-second resolution to enable us to compare network activity to ongoing behaviors widely utilized to assess the anxiety state of mice ^12,29^. This choice of one-second resolution balances temporal resolution and signal strength, as longer windows would be expected to yield stronger signals with lower temporal resolution. The 41 implanted male mice were exposed to at least 1 of the 3 experimental paradigms; 17 mice used to train the model were subjected to two paradigms (Supplemental Figure S2).

Our dCSFA-NMF model trained on the neural data acquired during the FLX paradigm successfully distinguished the low- and high-anxiety states in holdout subjects [saline and FLX treatment, respectively; Mann-Whitney Area under the curve of model performance (AUC) = 0.68 ± 0.01, where higher values indicated better model performance and 0.5 corresponds to models with no predictive utility; Fig. 1B, see also Supplemental Figure S3]; however, the model failed to distinguish low- and high-anxiety states when it was tested on data obtained from the other two paradigms (AUC = 0.49 ± 0.01 and 0.44 ± 0.01 for EPM and BOF, respectively). We also found that dCSFA-NMF models trained on the EPM or BOF assay similarly failed to distinguish the low- and high-anxiety states of the FLX assay. These analyses employed four-fold cross-validation with 3-7 holdout mice within assay per fold, and 9-26 holdout mice between assays per fold. A full discussion of the dCSFA-NMF model training procedure, hyperparameter selection, and rigorous validation strategies can be found in the Methods section.

After failing to discover a generalized internal state for anxiety solely using training data from one paradigm, we took inspiration from multi-task learning ^30^ and adapted dCSFA-NMF for training on multiple assays jointly (Fig. 1C, Supplemental Fig. S4). Specifically, the multi-assay dCSFA-NMF model utilized training data from all three contexts (FLX, EPM, BOF) to discover an electome network that was shared between them. Though we successfully discovered such a shared electome network, we also found that small permutations of the animal assignments between training and validation data groups yielded electome networks composed of different LFP spectral features (Fig. 1D), while remaining predictive of the high- vs. low-anxiety states.

To address this lack of stability, we developed and employed a cosine similarity-based metric for evaluating network stability across multiple training permutations. For this metric, a low cosine distance reflects greater electome network consistency. We then systematically increased the number of supervised networks in our dCSFA-NMF model, utilizing all supervised networks in a joint prediction logistic regression framework, and quantified the stability of the resultant electome networks. With this approach, we found that a model trained with three supervised electome networks optimally balanced simplicity and electome network stability across multiple runs as evaluated by cross-validation-over-subjects on training data (Supplemental Figure S5). For this multi-network model, higher activity was mapped to the high-anxiety contexts across all three assays. This multi-network model also encoded the high-anxiety context across all three assays in 20 newly implanted C57 mice that had not been used to train the model (AUC = 0.59 ± 0.04, 0.76 ± 0.03, and 0.84 ± 0.03 for FLX, EPM, and BOF, respectively; Fig. 1E). Although many individual brain regions/pairs of regions independently encoded one or two paradigms, no single region or pairs of regions encoded the anxiety state shared by all three paradigms (Fig. 1F, Supplemental Figure S6). Thus, our findings argued that the convergent anxiety brain state was encoded at the multi-region level. As expected, increasing the window length increases classifier discrimination (see Supplemental Figure S7).

### Two electome networks independently encode the anxious internal state

The three supervised networks that jointly predicted the anxiety state contributed 26%, 73%, and 1% of the joint logistic regression model prediction probability, averaged across assays (referred to as Electome Networks 1, 2 and 3, respectively; see Fig. 2D). Electome Network 1 was comprised of prominent beta (14–30Hz) and gamma oscillations (30–55Hz) that led from VTA, Amy, and MD, and converged in IL and NAc. Electome Network 2 was comprised of prominent beta and gamma oscillations that led from VHip, relayed through MD, and converged in the Amy. Electome Network 3 was represented by synchronized theta oscillations (4–11Hz) across many of the regions we probed (Fig. 2A-C, see also Supplemental Figures S8-10). Thus, the three electome networks were each represented by distinct ensembles of LFP activity.

**Figure 2:**
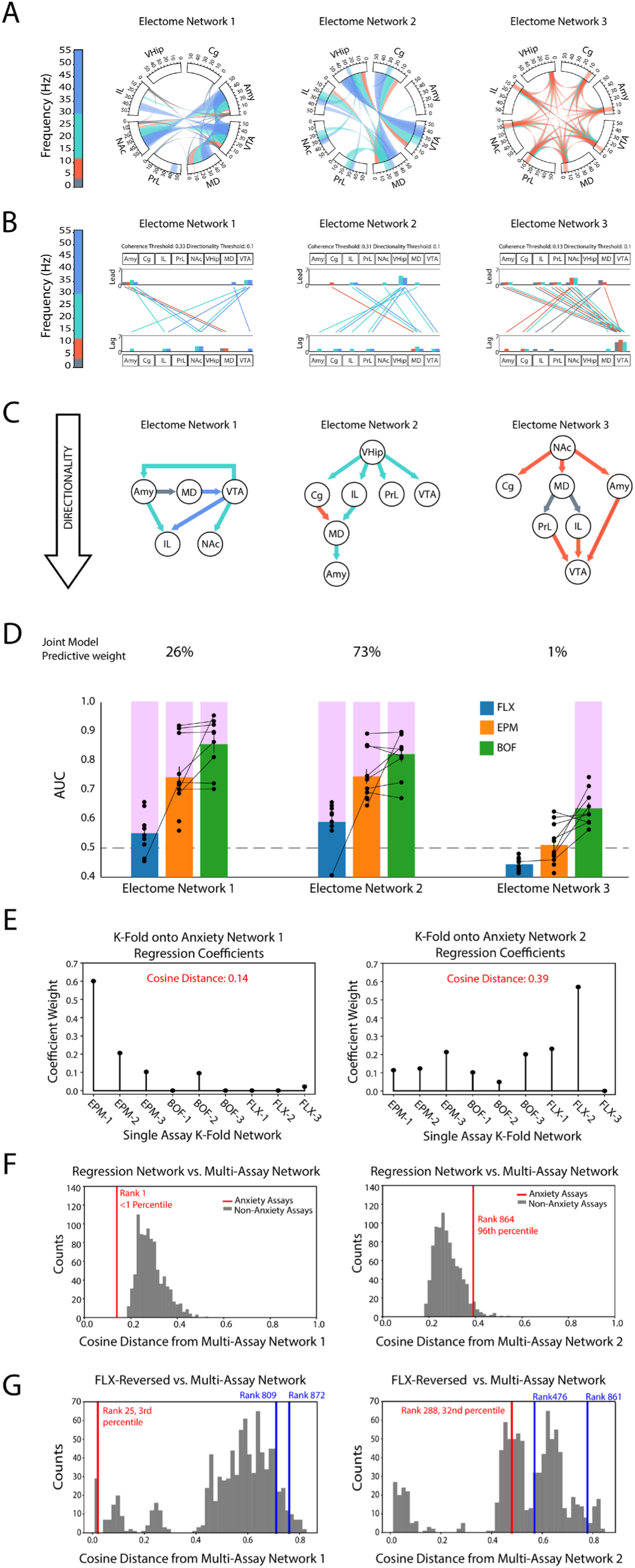
Individual electome networks within the multi-assay anxiety model independently encode distinct anxiety paradigms. **A)** Power, coherence, and directionality measures that comprise each electome network. Brain regions and frequency bands ranging from 1–56 Hz are shown around the rim of the circular plot. Power features are depicted as bands within the rim of the plot, and cross-spectral coherence measures are depicted by the lines connecting the brain regions through the center of the circle. The top 15% of components for each electome network are shown. **B)** Granger causality was used to quantify directionality for the coherence measures shown in A. Prominent directionality features were found in multiple bands coded by color. Histograms quantify the number of lead and lagging circuit interactions for each brain region. **C)** Schematic of directionality for each of the three electome networks. Arrows are colored to represent the dominant frequency of directionality (see color bar in panels A or B). **D)** Independent predictive performance of each supervised network across each anxiety assay. Mean contribution towards the joint model logistic regression predictions is also shown as the weight listed above each network’s graph. Tests were performed using 17 holdout mice, and networks that showed significant encoding are highlighted in pink (one-tailed unpaired t-test against a null of mean AUC of .5 at α = 0.05. Horizontal dashed line at AUC = 0.5 corresponds to models with no predictive utility. Data shown as mean±s.e.m. Solid lines indicate the data points from the same mouse. **E)** Coefficient weights for networks learned using individual assays (Fig. 1B and Supplemental Fig. S3) regressed onto Network 1 (left) or Network 2 (right). Cosine distances quantifying the difference between the single-assay regression network and the multi-assay networks are listed in red. **F)** Distribution of cosine distances between Network 1 (left) or Network 2 (right) and 900 regression networks generated from three assays of affective states orthogonal to anxiety. Each null distribution was compared to the cosine distances listed in E (shown as vertical red lines). **G)** Distribution of cosine distances between Network 1 (left) or Network 2 (right) and 900 multi-assay networks generated by randomly permuting the anxiety labels (low anxiety vs. high anxiety). The colored vertical lines indicate the cosine distances for the three networks learned using a model in which the fluoxetine and saline labels were reversed; the smallest cosine distances are noted in red. Given that Network 2 was not recovered under this reversed-label training, this network is henceforth referred to as *EN-Anxiety*.

Importantly, the electome networks included circuits previously implicated in aspects of anxiety-like behavior in the broader literature. For example, optogenetic stimulation of Amy → IL, a component of Electome Network 1, has been shown to increase anxiety-related behavior during the EPM and BOF in mice ^19^. Mouse studies have described increased activity in the VHip → prefrontal cortex circuit in the EPM and BOF ^12,13^, and causal stimulation of this circuit increases anxiety behavior ^22^. Increased coherence between Amy and VHip has been implicated in trait and state anxiety in human intracranial recording experiments ^8^ and in mediating EPM anxiety behavior in causal mouse experiments ^31^. Such circuits are prominently featured in Electome Network 2. Finally, increased IL activity, as featured in Electome Network 2 and 3, drives anxiety behaviors in mice in the EPM ^32^. Thus, many circuits previously shown to encode aspects of anxiety were featured in our discovered electome networks.

Though our goal was to discover at least one electome network that was shared across the three anxiety paradigms, our multi-supervised network learning strategy had the potential to discover three electome networks that each solely encoded one of the three assays. Thus, to ensure that we had indeed discovered an electome network that generalized across anxiety paradigms, we tested whether Electome Network 1, 2 or 3 encoded the anxious state in all three paradigms, again in the same 20 newly implanted mice mentioned above. Here we used one-tailed testing to determine whether higher activity in each network was observed in the high-anxiety context, as it had in our initial learned model. Electome Networks 1 and 2 both independently generalized to all three paradigms (P<0.05 for all comparisons against a null distribution, one-tailed Mann-Whitney U test), while Electome Network 3 only encoded the internal state induced by the BOF assay (P = 1, P=0.66; P <0.05 for FLX, EPM, and BOF, respectively; Fig. 2D). Additionally, since Electome Network 3 only contributed 1% to multi-network prediction, we limited our subsequent analysis to Electome Networks 1 and 2.

We next evaluated whether the architecture of Electome Networks 1 and 2 represented a brain state that was shared among the three assay contexts, or whether the networks simply represented the sum of the circuits that independently encoded each of the three assays. We quantified the extent to which our two multi-assay networks could be recovered from the individual assay models. Specifically, we started with the single assay-trained K-fold cross-validation models (from Fig. 1B), and we then selected the 3 supervised networks from the fold most predictive of each assay. This yielded 9 total supervised networks (3 for each assay), and we regressed these 9 networks onto Networks 1 and 2 from the multi-assay model (Fig. 2E). We assessed the quantitative fit between the multi-assay networks and the superimposed single-assay networks using the cosine distance. Here, a lower cosine distance indicated that a multi-assay network was similar to the superimposed single-assay networks. To generate a null distribution for statistical comparisons, we used three additional data sets for which there was no clear anxiety context (i.e., LFP recordings from social interaction test, sucrose countdown reward assay, and baseline sleep-wake recordings). We trained 10 single-assay models where the context labels (context 1 vs. context 2) were randomly assigned, generating 3 random supervised networks for each of the three data sets across 10 runs. We then randomly selected 9 networks from these 90 learned networks (3 networks from each of the data sets) and regressed them onto Networks 1 and 2. This pseudorandom selection procedure was repeated 900 times, yielding a null distribution consisting of 900 cosine distances between the multi-assay networks and the unrelated single-assay networks (Fig. 2F). A cosine distance less than 95% of this null distribution is equivalent to P<0.05, as the expected recovery is one-tailed.

The cosine distance between Electome Network 1 and the composite regression of the 9 supervised models (3 for each of the individual anxiety assays) was 0.14. This distance was less than 99% of the null distribution (Fig. 2F, left), providing statistical evidence that Network 1 may indeed represent a superposition of the single-assay networks. On the other hand, the cosine distance between Electome Network 2 and the composite regression of the 9 supervised models was 0.39. This distance was less than only 4% of the null distribution (Fig. 2F, right), providing evidence that Electome Network 2 was not a superposition of the single-assay networks.

Since we had previously found that the single-assay networks trained for the EPM assay generalized to the BOF assay and vice versa, our initial statistical approach did not allow us to disambiguate as to whether Network 1 represented a superposition of distinct individual assay networks or if it was strongly influenced by a single network shared between the EPM and BOF assays. Thus, to investigate this further, we retrained a new multi-assay model in which only the FLX and saline labels were reversed in the training set (FLX-reversed) while the EPM and BOF labels were not reversed. Based on our prior results, we expected that: (1) we would still recover Network 1 as it is primarily based on EPM and BOF; (2) this new FLX-reversed multi-assay network would classify FLX as the saline context and saline as the FLX context in holdout animals; and (3) we would fail to recover Network 2 since our initial analysis suggested it was dependent on all three assays.

All three supervised networks within the FLX-reversed multi-assay model generalized to holdout mice, predicting home cage vs. EPM, home cage vs. BOF, and reversed fluoxetine vs. reversed saline conditions. When we evaluated whether this procedure recovered the original Electome Network 1, again using the cosine distance, we found that the supervised FLX-reversed networks showed cosine distances of 0.71, 0.76, and 0.02 from Network 1 (Fig. 2G, left). To evaluate the significance of smallest network distance, we trained 900 anxiety-null multi-assay models for which the labels (low anxiety vs. high anxiety) for all three assays were randomly permuted. Then, we constructed a null distribution by choosing the supervised network with the smallest cosine distance from Network 1 for each permutation. The lowest cosine distance between the three FLX-reversed multi-assay networks and Network 1 (i.e., 0.02) was lower than nearly all the values in the null distribution (P=0.03; Fig. 2G, left). Thus, Network 1 was still recovered under FLX-reversed training. However, when we repeated this approach using Network 2, its lowest cosine distance with one of the three networks from the FLX-reversed multi assay model was 0.48. This cosine distance was indistinguishable from the null distribution (P=0.32; Fig. 2G, right), meaning that Network 2 was not recovered under FLX-reversed training. Taken together, these results established that Network 1 represented a network that was likely shared between the EPM and BOF assays, with a small input from the FLX assay. In contrast, Electome Network 2 represented an internal state that was shared across all three assays. Thus, we focused our subsequent analysis on Electome Network 2 (henceforth referred to as *Electome Network-Anxiety* or *EN-Anxiety*).

Finally, we also verified that *EN-Anxiety* generalized to female mice in the EPM assay (AUC=0.64±0.04, U=5, P= 0.0012, one-tailed paired Wilcoxon signed rank test; N=13 female mice) and the BOF assay using one-second windows (AUC=0.56±0.03, U=8, P= 0.048, one-tailed paired Wilcoxon signed rank test; N=9 female mice). As expected, encoding strength of these two assays increased with larger window sizes (Supplemental Figure S7). However, *EN-Anxiety* failed to generalize to female mice in the FLX assay (AUC=0.46±0.03, U=27, P= 0.90, one-tailed paired Wilcoxon signed rank test; N=8 female mice). When we explored this lack of generalization further, we found that treatment with fluoxetine (10mg/kg, i.p.) did not increase behavioral inhibition or heart rate in female mice (Supplemental Figure S11). Thus, the effects of our FLX assay failed to meet our operational definition of anxiety in female mice. Taken together, these results showed that *EN-Anxiety* selectively generalized to the two assays that successfully induced anxiety in female mice. Notably, we used a one-tailed statistical test for this analysis and our subsequent network validation testing, since no change or a decrease in network activity in the high-anxiety setting both equally reflected a failure of our learned *EN-Anxiety* to generalize to female mice. This approach also allowed us to limit the number of mice utilized for each assay, ultimately enabling us to probe network activity across a much broader range of anxiety-related and control contexts (see Methods – Statistical Analysis Philosophy).

### EN-Anxiety activity encodes features of anxiety-related paradigms

We further validated *EN-Anxiety* by examining network activity dynamics during various anxiogenic events, both within the training assays (i.e., FLX, EPM, and BOF) and in new experimental contexts that controlled for confounding emotional states. All analyses were performed on subjects that were not used in model training.

Within the FLX-training assay, we observed that the activity of *EN-Anxiety* decreased over time across the neural recording period in the saline- and FLX-treated mice (F_59,472_=2.88, P=0.038 for time effect for *EN-Anxiety*, within-subject two-way ANOVA with correction). As expected, given the AUC values we reported for the FLX assay in these same animals (see Fig. 1E), the activity of *EN-Anxiety* was higher in the fluoxetine-treated group using the same data as above for a confirmatory comparison (F_1,8_=14.81, P=0.005 for the treatment effect of the ANOVA). Nevertheless, no differences in the time effect were observed between the treatments (F_59,472_=1.49, P=0.22 for treatment × time interaction, two-way ANOVA with multiplicity correction; see Fig. 3A). Thus, following handling and the experimental injection, activity in *EN-Anxiety* decreased the longer mice they were in their home cage. These findings provided additional evidence that *EN-Anxiety* tracked the internal anxiety state of the mice.

**Figure 3:**
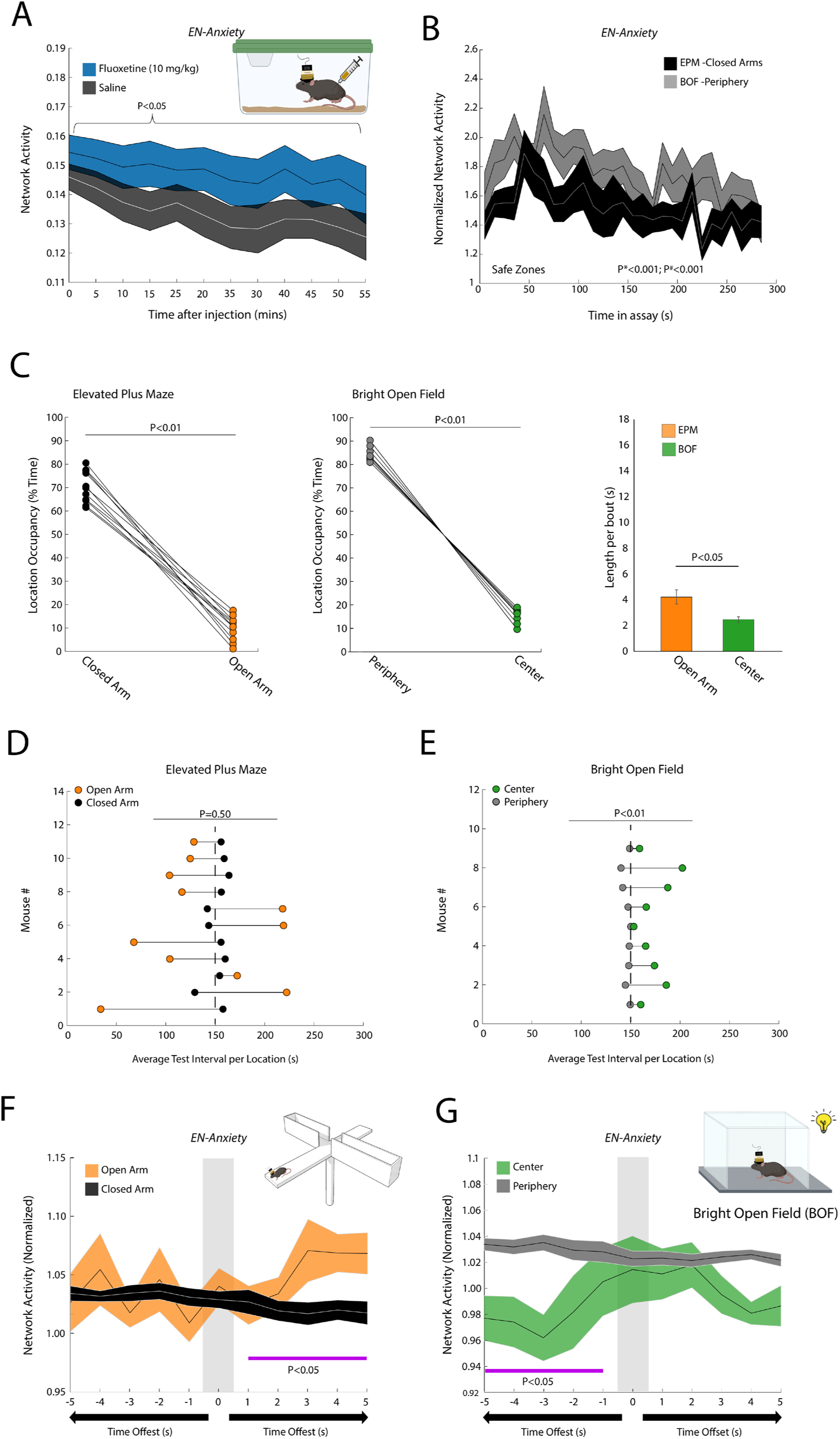
*EN-Anxiety* activity encodes features of anxiety-related paradigms. **A)** *EN-Anxiety* activity dynamics during fluoxetine assay. Data is plotted across 5-minute windows for *EN-Anxiety*, whose activity decreases over time following saline and fluoxetine treatment (N = 9 mice, P<0.05 for time effect, within-effects two-way ANOVA). **B)** Comparison of *EN-Anxiety* activity in safe zones of the EPM (closed arm) and BOF (periphery) over the duration of the assays. Time (P*) and assay (P^#^) effects were determined using an ANCOVA (see methods). Data was plotted with a 10s sliding window and shown normalized to network activity observed in the home cage. **C)** Mice showed avoidance of the ‘anxiogenic’ zones of the EPM (left, P<0.01, one-tailed paired t-test) and BOF (middle, P<0.01, one-tailed paired t-test). Bout length of mice in the avoidance zones for EPM and BOF (right; P<0.05 using one-tailed unpaired t-test). **D-E)** Average test interval per location/mouse in safe and (see methods) anxiogenic zones in D) EPM and E) BOF assays. The vertical dashed line at 150s divides the first and second halves of each assay. Note that mice showed greater occupancy of the center in the second half of the BOF, compared to the first half. **F)** *EN-Anxiety* activity dynamics relative to arm locations in the EPM assay. Gray highlights 1-second windows when the animals are in the open or closed arms. Neural activity preceding and following these timepoints is shown, normalized to the mean activity observed across the assay. The purple line highlights temporal intervals with significantly different *EN-Anxiety* activity, determined using a one-tailed Wilcoxon signed rank test (N = 11 mice). **G)** Same as F, except data is shown for the BOF assay (N=9 mice). Data (except in D-E) plotted as mean±s.e.m.

To further explore whether network activity decreased over time following exposure to other anxiogenic stimuli, we also analyzed network activity in the EPM and BOF, relative to that observed in the home cage. We focused on periods when mice were in the closed arms of the EPM or the periphery of the BOF, since these are considered safe zones. This approach also enabled us to control for changes in network activity that may be location specific. Activity in *EN-Anxiety* increased sharply after the mice were first placed in the behavioral areas, then decreased over time across the remainder of the 5-minute testing session (Fig. 3B; T=-11.116, P<0.001 for time effect for *EN-Anxiety*, ANCOVA; task activity is depicted normalized to home cage activity). Thus, network activity in both assays paralleled the response we observed in the FLX assay. Next, we tested whether network activity was behaviorally relevant. Specifically, since we found that the BOF induced higher network activity than the EPM (Fig. 3B; T=13.248, P<0.001 for assay effect, ANCOVA of assay and time), we analyzed the behavioral profiles of mice in both assays to determine whether the BOF induced greater anxiety-related behavioral avoidance. After verifying that the mice spent substantially more time in the safe zones vs. avoidance zones in both assays (T_10_=19.9; P<10^-8^ and T_8_=29.8; P<10^-8^, for EPM and BOF, respectively, one-tailed paired t-test, Fig. 3C, left and middle), we quantified the bout length when animals occupied the avoidance zones. The bout length in the center zone of the BOF was significantly shorter than in the open arm for the EPM (T_18_= 2.6; P=0.009, one-tailed unpaired t-test, Fig. 3C, right), demonstrating that mice exposed to the BOF showed higher anxiety-related avoidance. Mice avoided the center of the BOF more during the first half of the assay, compared to the second half (T_8_=3.97 and P=0.004, two-tailed paired t-test; see Fig. 3E). In contrast, no such behavioral pattern was observed in the EPM (Fig. 3D); mice showed large variability when they occupied the open arms (T_10_=0.70 and P=0.50, two-tailed paired t-test). These results suggested that mice were most likely to exhibit behavioral inhibition/avoidance when *EN-Anxiety* activity was highest (e.g., first half of the BOF).

To directly explore the relationship between *EN-Anxiety* and avoidance behavior, we tested whether network activity encoded occupancy of the avoidance zone on a moment-to-moment basis within the assays. Specifically, we reasoned that three distinct patterns of anxiety could intersect with behavior: 1) mice might show higher anxiety when they occupy the avoidance zones of the assay, which could be indicated by higher network activity in open arm/center vs. closed arm/surround; 2) the avoidance zone of the assays might induce a feeling of anxiety that increases over time irrespective of the animal’s future location, observed as higher network activity seconds after mice are located in the open arm/center vs. seconds after they are located in the closed arm/surround; and 3) high anxiety might preclude mice from occupying the avoidance zones ^31^, such that higher network activity would be observed if mice occupy the closed arm/surround in the future vs. when mice occupy the open arm/center. Thus, we determined whether *EN-Anxiety* showed activity consistent with any of these 3 patterns. Importantly, though all data recorded during the EPM and BOF assays were used to discover our putative anxiety networks, the moment-by-moment locations of the mice in the EPM and BOF (avoidance zone vs. safe zone) were not. Thus, this analysis tests the relationship between *EN-Anxiety* activity and behavior based on information that was not used during model training.

We first isolated all the one-second segments when mice occupied the open or closed arm of the EPM. We also isolated neural activity up to five seconds prior to and up to five seconds following these timepoints ^15,31^. *EN-Anxiety* failed to encode whether mice were in the open vs. closed arms of the EPM (U=41 and P=0.26, one-tailed paired Wilcoxon signed rank test; Fig. 3F); this failed to support the first pattern of anxiety-related network activity listed above. On the other hand, we found that *EN-Anxiety* activity was higher in the five seconds interval following the open arm location of mice (U=61 and P=0.0049, one-tailed paired Wilcoxon signed rank test), regardless of whether they returned to the closed arm during this period. Thus, an increase in *EN-Anxiety* activity was induced by the avoidance zone of the assay, providing support for the second pattern listed above. No difference in network activity was observed in the five-second interval preceding the open arm location (regardless of the location of the mouse during this interval), compared to activity preceding the closed arm of the EPM (U=27 and P=0.32, one-tailed paired Wilcoxon signed rank test; Fig. 3G). This failed to support the third pattern for which high network activity might preclude entrance into avoidance zones.

*EN-Anxiety* also failed to encode whether mice were in the center or periphery of the BOF (U=20 and P=0.63, one-tailed paired Wilcoxon signed rank test), again showing a lack of support for the first pattern. However, when mice occupied the center of the BOF, the network showed lower activity within the preceding five seconds, compared to the five seconds preceding when mice occupied the periphery (U=8 and P=0.049, one-tailed paired Wilcoxon signed rank test, Fig. 3G). Together, this finding showed that high activity in *EN-Anxiety* encoded that mice would be in the safe zone of the BOF in the future, thus supporting the third pattern. No increases in *EN-Anxiety* were observed in the five-second interval following the center location compared to activity when mice were in the periphery in the BOF (U=13 and P=0.88, one-tailed paired Wilcoxon signed rank test). Overall, these results showed that *EN-Anxiety* activity was increased by the open arms of the EPM, while high activity in *EN-Anxiety* encoded avoidance of the center zone in the BOF. This latter pattern of activity was consistent with our observation that mice occupied the center zone less during the first half of the BOF when *EN-Anxiety* activity was highest. Thus, *EN-Anxiety* was behaviorally relevant and supported two out of three of our patterns of anxiety-related network activity, though the pattern for which aspects of anxiety behavior were encoded varied between the two assays.

### EN-Anxiety activity does not encode arousal or reward

As arousal is a component of our operational definition of anxiety, our training approach could plausibly discover networks that reflect an arousal state rather than anxiety. To explore this possibility, we quantified *EN-Anxiety* activity during distinct sleep-wake arousal states (Fig. 4A). Specifically, we recorded animals for 12 hours and identified intervals when animals were in the middle of long bouts of wakefulness or in rapid-eye-movement (REM) sleep ^33^ (Fig. 4B). Though REM sleep is characterized by fast brain activity, the body shows broad atonia, and animals are less responsive to external stimuli. As such, it is considered a lower arousal state than wakefulness. Muscle activity was higher during the awake period compared to the REM sleep intervals (U=36, P=0.0039, one-tailed paired Wilcoxon signed rank test; N=8 mice; Fig. 4C). This was consistent with our sleep-wake classification approach^33^ and reflected the higher arousal state that characterizes wakefulness.

**Figure 4:**
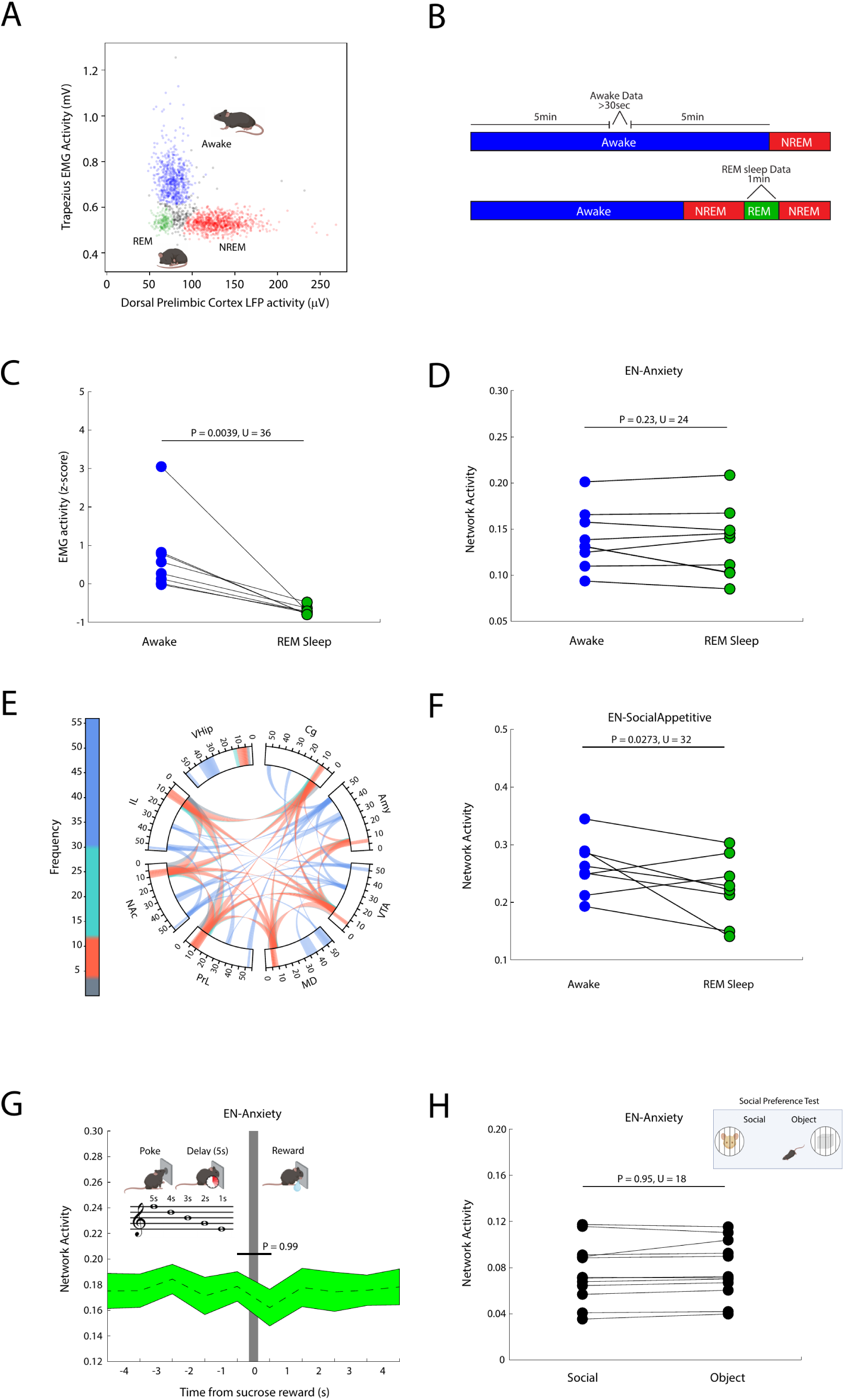
*EN-Anxiety* does not encode orthogonal affective states. **A)** Depiction of clustering approach utilized to classify sleep-wake states (see Methods for details; unlabeled windows are shown in black, awake windows are in blue, REM windows are in green, and NREM windows are in red.). **B)** Selection criteria for awake and REM sleep periods for analysis. Awake intervals were defined by at least 30 seconds of awake-labeled data with at least 5 minutes of >95% awake labels on both sides of the data segment (top). REM sleep intervals were defined by at least 60 consecutive seconds that were labeled as REM sleep (bottom). **C)** Average electromyographic activity during selected awake and REM sleep intervals. **D)** *EN-Anxiety* network activity during awake vs. REM sleep periods (P>0.05 using one-tailed paired Wilcoxon signed rank test). **E)** *EN-SocialAppetitive* network previously discovered to encode a social appetitive brain state^26^. Brain regions and frequency bands ranging from 1–56 Hz are shown around the rim of the plot. Power features are depicted as bands within the rim of the plot, and cross-spectral coherence measures are depicted by the lines connecting the brain regions through the center of the circle. The top 15% of components is shown. **F)** *EN-SocialAppetitive* network activity during awake vs. REM sleep periods (P<0.05, one-tailed paired Wilcoxon signed rank test). **G)** Mice were trained to maintain a nose poke for 5 consecutive seconds. A sucrose reward was delivered at time zero, highlighted by gray. *EN-Anxiety* activity was compared before and after sucrose delivery using a one-tailed paired Wilcoxon signed rank test (N=9 mice). Data is shown as mean±s.e.m. **H)** *EN-Anxiety* activity was quantified while mice engaged with an object or a social stimulus mouse during a free interaction assay and compared using a one-tailed sign-rank test (N=12 mice). All analyses were performed in mice that were not used to learn the multi-assay anxiety model.

We reasoned that *EN-Anxiety* activity would be higher during wakefulness compared to REM if the network directly encoded arousal, rather than anxiety more broadly. Though *EN-Anxiety* activity was not significantly higher during wakefulness (U=24, P=0.23, one-tailed paired Wilcoxon signed rank test; Fig. 4D), such a finding might reflect an inability of networks discovered using our approach to encode differences between wakefulness and REM. To exclude such a possibility, we tested whether one of our previous electome networks that encodes social appetitive behavior (*EN-SocialAppetitive*, Fig. 4E, see also Supplemental Fig. S12)^26^ would show higher activity during wakefulness, compared to sleep. Importantly, this network is composed of the same brain regions as *EN-Anxiety*. When we projected the data from the wakefulness and REM sleep intervals into this electome network, we found that network activity was indeed significantly higher during wakefulness vs. REM sleep (U=32, P=0.0273, one-tailed paired Wilcoxon signed rank test, Fig. 4F). Thus, changes in network activity between wakefulness and REM sleep could indeed be captured by a network composed of the same brain regions as *EN-Anxiety*, but not by *EN-Anxiety* itself. Together, these results showed that *EN-Anxiety* did not simply encode arousal.

We next tested whether increases in *EN-Anxiety* network activity were specific to anxiety, using data acquired from two independent assays that are thought to increase reward but not anxiety. During these assays, data was collected from the same brain regions used initially for model training for *EN-Anxiety*, and LFP activity was projected into the previously learned multi-assay trained model to calculate the activity of *EN-Anxiety* for each second. In the first assay, nine mice were trained to maintain a nose poke for 5 seconds (Supplemental Fig. S13). Tones of decreasing pitch were played throughout the 5-second trial, and a 5µL sucrose reward was delivered at the end if mice remained in the port for the entire 5 seconds. When we tested whether reward delivery increases *EN-Anxiety* activity, we failed to identify a significant increase in *EN-Anxiety* after sucrose reward delivery (U=41, P=0.99, one-tailed paired Wilcoxon signed rank test; N=9 mice, Fig. 4G). Next, we quantified *EN-Anxiety* activity during a classic social preference assay where mice freely explore an inanimate object and a novel social stimulus mouse housed at the two ends of a chamber. In this assay, the social stimulus mouse is considered both arousing and rewarding, as experimental mice generally choose to spend more time with the other mouse than the object, and social encounters activate reward circuitry^34^. When we projected

LFP data collected in a prior study^26^ into *EN-Anxiety*, we failed to discover increases in *EN-Anxiety* during interactions with the stimulus mouse, compared to the inanimate object (U=47, P=0.74, one-tailed paired Wilcoxon signed rank test, N=12 mice, Fig. 4H). Together, these results demonstrate that *EN-Anxiety* fails to encode arousal and reward, as no increase in network activity is observed during higher arousal states or during innate and trained-reward states.

### EN-Anxiety activity encodes the internal state induced by additional anxiety paradigms

We further probed whether *EN-Anxiety* encodes a robust anxiety-related brain state using additional paradigms that induce anxiety in a manner that is distinct from our initial assays. Again, all analyses were performed on new mice that were not used during model training. We first examined network activity during direct optogenetic stimulation of neurons in the VHip. This region has been causally implicated in anxiety in rodents ^35^, and it was a critical upstream node in *EN-Anxiety*. Moreover, we selectively stimulated the subset of neurons that projected to lateral hypothalamus (LH) since the VHip → LH circuit had been shown to drive anxiety-related avoidance in the EPM and BOF ^20^. Mice were infected with an adenoassociated virus (AAV) to express Channelrhodopsin-2 using (ChR2) in the VHip and implanted with microwires to target the same regions utilized to learn *EN-Anxiety*. A microwire and optic stimulating fiber were also implanted in LH, concurrently (Fig. 5A).

**Figure 5:**
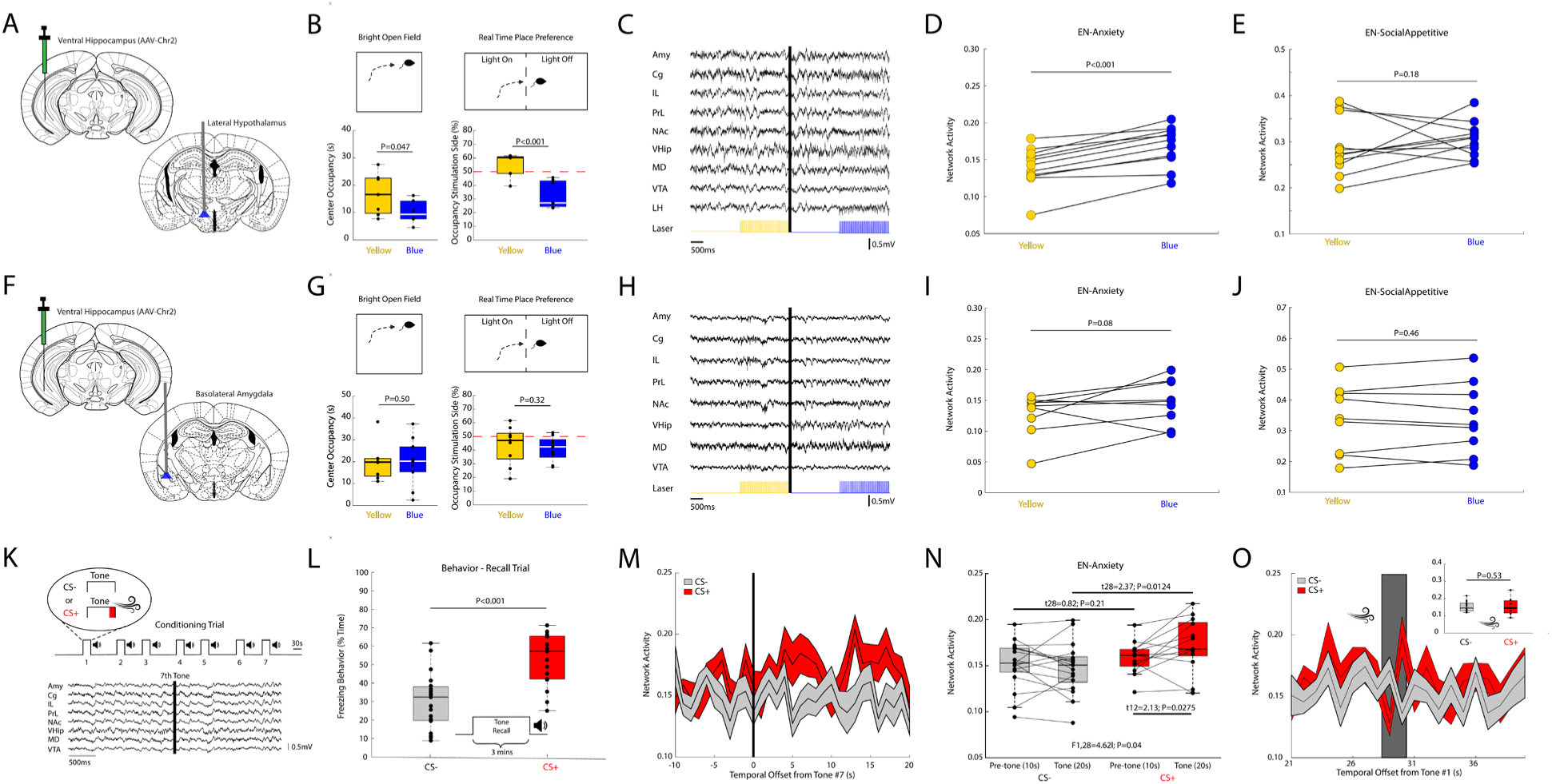
*EN-Anxiety* activity encodes distinct anxiety paradigms. **A)** Mice were infected with ChR2 in ventral hippocampus (VHip) and a stimulating fiber was implanted in lateral hypothalamus (LH). **B)** Mice (N=13) were stimulated with blue (473nm, 20Hz, 5mW, 5ms pulses) or yellow light (593.5nm, 20Hz, 5mW, 5ms pulses) during testing in the BOF (left), or one side of a real-time place preference assay (right). Comparisons of yellow light and blue light stimulation were run with a one-tailed t-test. **C)** Mice were infected with ChR2 in VHip and an optrode was implanted in LH. Multiwire electrodes were also implanted to target the 8 brain regions utilized to learn the multi-assay anxiety network (N=11 mice). Neural activity recorded during optogenetic stimulation of VHip terminals in LH with yellow (left) or blue light (right). Blue light stimulation of VHip induced activity in LH (and remotely) while yellow light stimulation did not. **D)** *EN-Anxiety* showed an increase in activity with blue vs. yellow light stimulation (P<0.001, one-tailed Wilcoxon signed rank test). **E)** In contrast, blue light stimulation failed to increase *EN-SocialAppetitive* activity (P=0.18, one-tailed Wilcoxon signed rank test). Mice (N=18) were infected with ChR2 in VHip and a stimulating fiber was implanted in basolateral amygdala (Amy). Mice were stimulated with blue or yellow light during testing in the BOF (left), or one side of a real time place preference assay (right). **H)** Mice were infected with ChR2 in VHip and an optrode in Amy. Multiwire electrodes were also implanted to target the 8 brain regions utilized to learn the multi-assay anxiety network (N=11 mice). Neural activity recorded during optogenetic stimulation of VHip terminals in Amy with yellow (left) or blue light (right). **I)** Blue light stimulation had no impact on *EN-Anxiety* or **J)** *EN-SocialAppetitive* activity. **K)** Behavioral paradigm utilized to induce fear conditioning. Conditioned mice (CS+; N=17) received an air puff at the end of each tone presentation, while non-conditioned mice (CS-; N=20) did not (top). Neural activity was recorded in both groups throughout tone presentation (bottom). **L)** Freezing behavior in CS- and CS+ mice one to two days after exposure to the conditioning paradigm. **M)** *EN-Anxiety* activity relative to the 7^th^ tone, tone onset denoted by the vertical black line (N=13 CS+ and 17 CS-mice; data shown as mean±s.e.m.). **N)** Mean *EN-Anxiety* activity within the interval prior to ‘pre-tone (10s)’] or during [’tone (20s)’] the presentation of the 7^th^ conditioning tone. Presentation of the 7^th^ tone increased network activity in the CS+ mice relative to CS-mice ([two-way repeated measures ANOVA, followed by a post-hoc one-tailed t-test). **O)** Mean activity of *EN-Anxiety* in response to the first air puff (stimulus presentation shown by dark grey vertical bar; N=7 CS+ and 15 CS-mice; data shown as mean±s.e.m.). Data was analyzed using a one-tailed Mann-Whitney rank sum test. Mean network activity for the two groups is shown in the inset. For the box and whisker plots, the central mark is the median, the edges of the box are the 25th and 75th percentiles, the whiskers extend to the most extreme datapoints the algorithm does not consider to be outliers.

First, we verified that VHip → LH stimulation recapitulated the anxiety/avoidance behaviors described in the prior work (t_16_=1.83, P=0.047; t_11_=4.09, P=8.9x10^-4^ for BOF center occupancy and real-time stimulation side occupancy, respectively, for comparisons of yellow and blue light stimulation; one-tailed t-tests, reflecting the *a priori* hypothesis that circuit stimulation would increase anxiety behavior; see Fig. 5B)^20^. Next, we stimulated the implanted mice with blue light to activate ChR2, or yellow light as a negative control, while neural activity was recorded in their home cage. As expected, blue light induced local and remote LFP activity, while yellow light did not (Fig. 5C). When we projected neural activity recorded during these stimulations into our learned multi-assay trained model, we found that VHip → LH stimulation increased *EN-Anxiety* activity (U=0, P<0.001, one-tailed paired Wilcoxon signed rank test, Fig. 5D). Stimulation of this circuit failed to increase *EN-SocialAppetitive* activity in the mice (U=22, P=0.18, one-tailed paired Wilcoxon signed rank test, Fig. 5E), demonstrating the circuit’s functional selectivity in driving activity in specific electome networks. In control studies, stimulation of a VHip → basolateral amygdala circuit (Fig. 5F) did not induce anxiety (t16=0.01, P=0.50; t_16_=0.48, P=0.32 for BOF center occupancy and real-time stimulation side occupancy, respectively; one-tailed t-tests for comparisons of yellow light and blue light stimulation; see Fig. 5G) and failed to significantly increase *EN-Anxiety* (U=35, P=0.08, one-tailed paired Wilcoxon signed rank test, Fig. 5H-I). This stimulation failed to increase *EN-SocialAppetitive* as well (U=21, P=0.46, one-tailed paired Wilcoxon signed rank test, Fig. 5J). Together, these data further validate *EN-Anxiety* as a network-level code that is specific for anxiety.

We subsequently examined whether *EN-Anxiety* encodes the internal state induced by fear conditioning. In this classic paradigm, mice are exposed to seven repeated auditory cues (conditioned stimuli, CS+), each paired to a foot shock (unconditioned stimulus, US). On a subsequent recall session, conditioned mice exposed to the auditory cue in the absence of the foot shock typically exhibit a freezing response. For our conditioning paradigm, we substituted the foot shock with a high-pressure air puff during conditioning (Fig. 5K). This enabled us to minimize electrical noise during LFP recording. Mice exposed to our air-puff stimulus exhibited increased freezing behaviors during the recall session on a subsequent day, compared to controls (CS-) (U=439, P=2.1×10^-4^, one-tailed Mann-Whitney U test; Fig. 5L). This confirmed that mice exhibited vigilance patterns that facilitated the association of a threat (US) with the context it was experienced (CS+) during the initial pairing session, meeting our operational definition of anxiety.

We then analyzed *EN-Anxiety* activity at the final stimulus of the aversive conditioning (i.e., the 7^th^ tone), since we reasoned that the six preceding air puffs were sufficient to induce vigilance in the mice. Specifically, we compared *EN-Anxiety* activity immediately prior to and during the 7^th^ tone between the CS+ and CS-mice. Here, we found a significant group by tone interaction (F_1,28_=4.62; P=0.04, two-way repeated measures ANOVA, N=13 CS+ and 17 CS-mice, respectively; Fig. 5M and 5N). Compared to the CS-control mice, *EN-Anxiety* activity was significantly elevated in CS+ mice during (t_28_=2.37, P=0.0124, one-tailed t-test, Fig. 5N), but not prior to (t_28_=0.82, P=0.21, one-tailed t-test, Fig. 5N), the presentation of the 7^th^ tone. Importantly, our post-hoc analysis found no difference in *EN-Anxiety* activity prior to or during the first tone exposure (i.e., preceding the first air puff in CS+ mice; F_1,1_=10^-4^ and P=0.99 for group effect, F_1,28_=0.55 and P=0.46 for tone effect, F_1,28_=0.56 and P=0.46 for group × tone interaction, two-way repeated measures ANOVA; see Supplemental Figure S14). As such, *EN-Anxiety* encodes an acute internal state generated by the presentation of a threat-paired stimulus.

Finally, having discovered that *EN-Anxiety* encoded threat presentation during hypervigilance, but not arousal or reward, we asked whether the network broadly encoded an acute negative experience. Specifically, while anxiety is a negative affective state, not all negative experiences produce anxiety. Thus, we further probed our fear conditioning paradigm data to test whether the *EN-Anxiety* increased acutely during an ongoing negative experience. We reasoned that prior to conditioning, the first air puff should immediately invoke negative affect, but not anxiety. As such, we compared *EN-Anxiety* activity in the CS+ mice while they experienced the first air puff to network activity in CS-mice, which did not receive an air puff. The air puff failed to acutely increase *EN-Anxiety* activity (U=80, P=0.53, one-tailed Mann-Whitney U test, Fig. 5O), which indicated that *EN-Anxiety* activity was indeed specific for anxiety rather than a generalized negative affective state.

### EN-Anxiety activity is altered in mouse models of mood disorders

Anxiety can be altered in mood disorders. Indeed, bipolar mania is characterized by impulsivity and risk taking^36^ (reflective of decreased anxiety processing), while major depressive disorder is highly co-morbid with high anxiety^37^. Mouse models of mood disorders have played a key role in investigating the mechanisms underlying these changes in anxiety. For these models, anxiety states in mice are generally inferred using assays such as the EPM. Yet, it is widely known that EPM behavior is shaped by other domains of affective function that are also altered in mood disorders (i.e., psychomotor drive, see Fig. 6A). Thus, illness-related changes across broader affective systems of mouse models can obscure the relationship between EPM behavior and the true anxiety state of mice. Having robustly established and validated *EN-Anxiety*, we reasoned anxiety levels could be directly assayed in mouse models of mood disorders by quantifying network activity. First, we tested whether a mouse model of mania would show lower *EN-Anxiety* activity in a context that should otherwise induce a high-anxiety state. We also tested whether two mouse models of depression would exhibit high *EN-Anxiety* activity in a context that would normally be marked by low anxiety.

**Figure 6:**
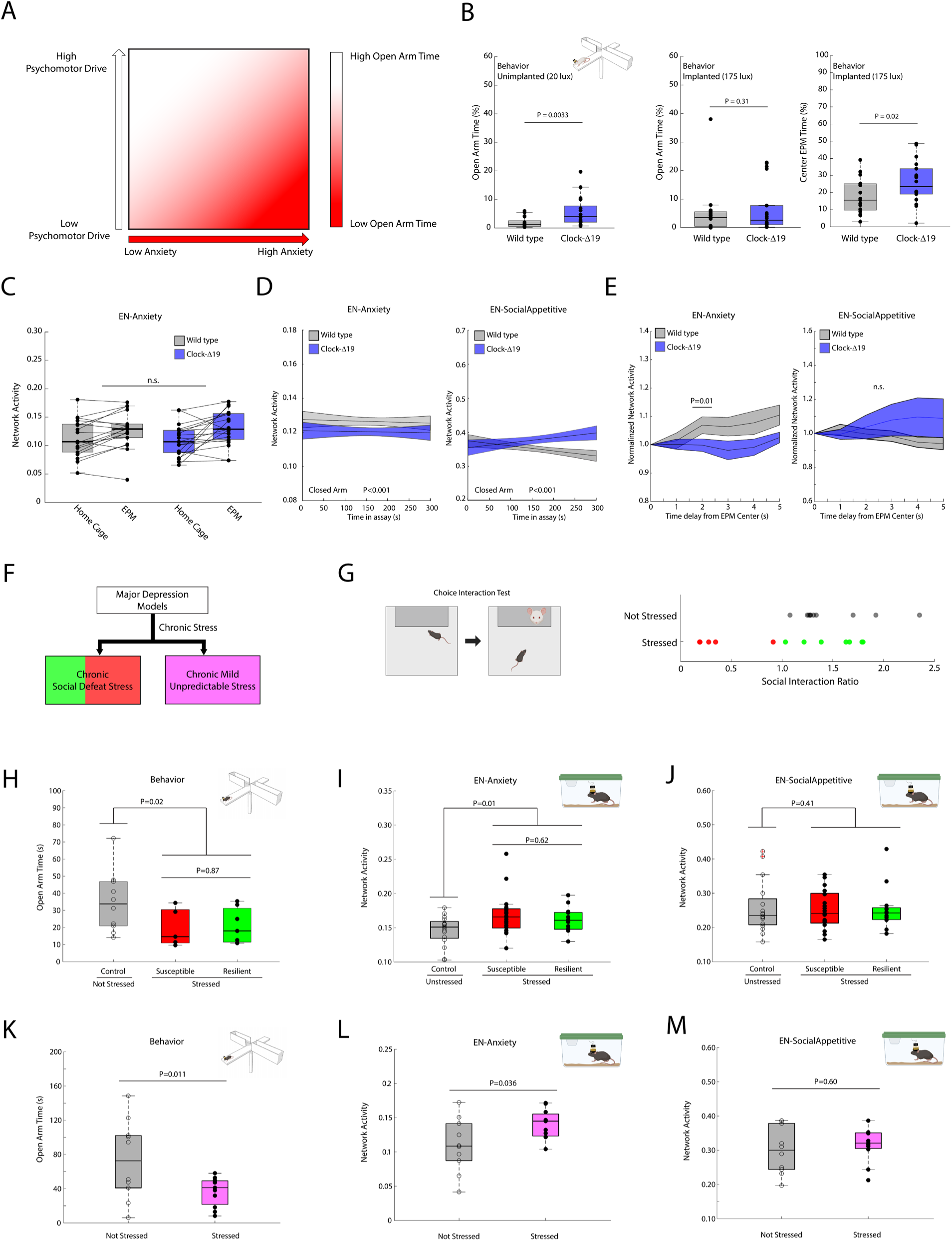
*EN-Anxiety* based assessment of anxiety function in mouse models of mood disorders. **A)** Schematic of how orthogonal affective states shape behavior in the EPM. **B)** EPM open arm exploration in unimplanted WT and *ClockΔ19* mice tested at 20 lux (N=17 mice/genotype, left), and implanted WT and *ClockΔ19* mice tested at 175 lux (N=17-18 mice/genotype, middle). Center occupancy time for the implanted mice is shown to the right. Data was compared using an unpaired one-tailed t-test. **C)** *EN-Anxiety* network activity while WT and *ClockΔ19* mice were in their home cage and in the EPM. Data was analyzed using a two-way repeated measures ANOVA. **D)** *EN-Anxiety* (left) and *EN-SocialAppetitive* (right) activity while WT and *ClockΔ19* mice were in the closed arm of the EPM. Genotype effects were determined using an ANCOVA (P<0.001 for both networks). Data visualized as mean±95%C.I. for network activity from each genotype independently fitted to a line. **E)** *EN-Anxiety* (left) and *EN-SocialAppetitive* (right) activity in response to occupancy of the center zone of the EPM. Data was compared across genotype using an unpaired one-tailed rank sum test, followed by an FDR correction for each offset. **F)** Distinct stress paradigms utilized to model depression in mice. Red and green divisions in the chronic social defeat stress paradigm represent the susceptible and resilient mice that result from the paradigm, respectively (left). Pink in the chronic mild unpredictable stress paradigm represents the stressed mice subjected to the paradigm (right). **G)** Schematic of choice interaction test utilized to quantify susceptibility to chronic social defeat stress (left), and resultant social interaction profiles of a population of stressed mice (right). Red circles denote susceptible mice (interaction ratio < 1), while green circles denote resilient mice (interaction ratio >= 1). Black circles denote non-stressed control mice. **H)** EPM open arm exploration in mice subjected to chronic social defeat stress (N=12 mice) and control mice (N=10 mice). Data was compared between stressed and non-stressed mice using a one-tailed t-test. Post-hoc testing between susceptible (N=5 mice) and resilient mice (N=7 mice) was performed using a two-tailed t-test. **I)** *EN-Anxiety* activity was quantified in the home cage and compared between chronic social defeat-stressed (N=34 mice) and non-stressed controls (N=16 mice) with a one-tailed Mann-Whitney rank-sum test. Post-hoc testing was compared between susceptible (N=21 mice) and resilient mice (N=13 mice) using a two-tailed Mann-Whitney rank-sum test. **J)** Comparison of *EN-SocialAppetitive* activity in the mice shown in G. Data was analyzed using a one-tailed rank-sum test. **K)** EPM open arm exploration in mice subjected to chronic mild unpredictable stress (N=11 mice) and control mice (N=11 mice). Data was compared using a one-tailed Welch’s t-test. **L)** *EN-Anxiety* activity was quantified in the home cage and compared across groups using a one-tailed Mann-Whitney rank-sum test. **M)** Comparison of *EN-SocialAppetitive* activity in the mice shown in J. For the box and whisker plots, the central mark is the median, the edges of the box are the 25th and 75th percentiles, the whiskers extend to the most extreme datapoints the algorithm does not consider to be outliers.

The *ClockΔ19* mouse line has been proposed as a model of bipolar mania ^38^. These mice have a point mutation in the circadian gene *Clock* and exhibit altered circadian rhythms, psychomotor hyperactivity, increased reward drive, and decreased avoidance in anxiety assays ^15,38,39^. Moreover, many cellular, neurophysiological, and behavioral alterations in these mutant mice are normalized by chronic lithium or valproic acid treatment ^38,40,41^, providing further validation for the *ClockΔ19* mouse as a model of bipolar mania. We reasoned that if *Clock* gene disruption altered anxiety, we would observe failure of *EN-Anxiety* to activate in response to EPM exposure, or an overall decrease in their *EN-Anxiety* activity. After confirming that *ClockΔ19* mice demonstrate increased open arm exploration in the EPM at 20 lux compared to wild-type littermate controls (t_32_=2.90, P = 0.0033, unpaired one-tailed t-test, Fig. 6B), we implanted male and female *ClockΔ19* mice on a Balb/c background (N=18 mice) and their wild-type littermate controls (N=17) with microwires targeting the same brain regions used to learn our electome network. We then quantified neural activity while mice were in the home cage and on the EPM. For these experiments, we used the same light intensity we had implemented in the C57 mice to learn and validate *EN-Anxiety* (i.e., 175 lux).

No differences in open arm exploration were observed across genotype at this higher light intensity (t33=0.51, P = 0.31, unpaired one-tailed t-test, Fig. 6B, middle). Moreover, exposure to the EPM increased *EN-Anxiety* activity in both genotypes, confirming that *EN-Anxiety* – discovered in C57 mice – generalized to a new mouse strain background (F_1,35_ = 14.21, P=6.05×10^-4^ for assay effect, two-way repeated-measures ANOVA for genotype × assay; Fig. 6C). Nevertheless, no significant interaction or genotype effect was observed (F_1,35_=0.73 and P=0.40 for genotype × assay interaction, F_1,35_=0.008 and P=0.93 for genotype effect; two-way repeated-measures ANOVA; Fig. 6C). Thus, gross *EN-Anxiety* function remained intact in *ClockΔ19* mice. This was consistent with the behavior displayed by the mutants, which showed strong avoidance of the open arm of the EPM (open arm occupancy = 5.8±1.3% and 5.2±2.1% in the unimplanted and implanted mice, respectively, Fig. 6B, middle). Indeed, nearly half (8 out of 18) of the implanted *ClockΔ19* mice occupied the open arm for less than 5 seconds.

Next, we wondered whether the *ClockΔ19* mice might exhibit lower *EN-Anxiety* network activity in the safe zone of the EPM, i.e., the closed arm. We indeed observed lower *EN-Anxiety* network activity in these mutant mice compared to their littermate controls (T=3.71, P<0.001 for genotype effect on *EN-Anxiety*, ANCOVA, Fig. 6D, left). The mutants also exhibited higher *EN-SocialAppetitive* network activity in the closed arms (T=4.05, P<0.001 for genotype effect on *EN-Anxiety*, ANCOVA, Fig. 6D, right), demonstrating that decreased network activity was specific to *EN-Anxiety*. Since these observations raised the possibility that the implanted *ClockΔ19* mice might experience decreased anxiety states in the safe zones compared to their wild type littermates, yet show similar open arm exploration, we wondered whether genotype differences were evident at the transition zone between the open and closed arms (i.e., the center zone of the EPM). Indeed, the implanted *ClockΔ19* mice exhibited greater occupancy of the center zone than their wild type littermates (t33=2.14; P=0.02, unpaired t-test; Fig. 6B, right).

Next, since occupancy of the open arm increased *EN-Anxiety* activity in C57 mice (Fig. 3F), and the center zone induced less avoidance in *ClockΔ19* mice compared to their wild type littermates, we hypothesized that *ClockΔ19* mice might also show reduced induction of *EN-Anxiety* activity following occupancy of the center zone. To probe this question, we isolated all 1-second periods when mice were in the center zone and determined network activity up to 5 seconds into the future, irrespective of the future location of the mice. This future network activity was normalized to the activity measured while mice were in the center zone, enabling us to compare the extent to which occupancy of the center zone impacted network activity across genotype. With this approach, we found that network activity was indeed lower in the mutants two seconds in the future (U=348, P=0.0099, unpaired one-tailed Wilcoxon rank sum test followed by false discovery rate correction, Fig. 6E, left), and tended to be lower 5 seconds into the future, though this difference did not survive correction for multiple comparisons (U =398, P=0.046, unpaired one-tailed Wilcoxon rank sum test followed by false discovery rate correction, Fig. 6E, left). No genotype differences were observed when we tested whether *ClockΔ19* disruption reduced *EN-SocialAppetitive* activity across these same time windows (P>0.05 for all comparisons using unpaired one-tailed Wilcoxon rank sum test followed by false discovery rate correction, Fig. 6E, right), demonstrating that the decreased network activity was specific to *EN-Anxiety*. Thus, *ClockΔ19* disruption decreased avoidance of the center zone of the EPM and suppressed the induction of *EN-Anxiety* in this otherwise high-anxiety context. Overall, these results show that a mouse model of bipolar mania exhibits decreased function of *EN-Anxiety* in a high-anxiety context, supporting the translational utility of this network.

Finally, we explored network activity in two mouse models of depression (Fig. 6F). In major depressive disorder, high anxiety is observed in contexts where it should typically be low/absent. Thus, we hypothesized that higher *EN-Anxiety* activity would be observed when the two mouse models of depression were in the low-anxiety context of their home cage. In the chronic social defeat stress paradigm, mice are repeatedly exposed to larger, aggressive mice. After 10 exposures, a subset of *susceptible* mice exhibits social avoidance, disrupted reward behavior, and anxiety-like behavior ^42,43^. Conversely, the other subset of *resilient* mice exhibits normal social and reward behavior ^42,43^ (Fig. 6F-G). Interestingly, despite the well-described differences in social appetitive behavior, prior work has reported increased open arm avoidance in the EPM in both the susceptible and resilient mice ^42,44^. Indeed, exposure to chronic social defeat stress increases open arm avoidance in the EPM for both unimplanted susceptible and resilient mice uniformly (t_20_=2.27, P=0.02, one-tailed Welch’s t-test for comparison between stressed and non-stressed mice; t_10_=0.16, P=0.87, two-tailed t-test for post-hoc comparison between susceptible and resilient mice; Fig. 6H). Therefore, we tested whether this stress paradigm also increased *EN-Anxiety* in both the susceptible and resilient groups compared to non-stressed controls. Again, since we hypothesized that stress exposure induces anxiety – and therefore higher *EN-Anxiety* activity – in contexts where anxiety is not observed under normal conditions, we quantified network activity from stress and non-stressed mice while they were in their home cage. Here, we found significantly higher *EN-Anxiety* activity in the stressed mice (U=151, P<0.01, one-tailed Mann-Whitney U test, Fig. 6I). Moreover, no difference in *EN-Anxiety* activity was observed between susceptible and resilient mice (U=382 and P=0.62, two-tailed Mann-Whitney U test for post-hoc analysis, Fig. 6I). Finally, no group differences in *EN-SocialAppetitive* activity were observed, demonstrating that the stress-induced changes in network activity were specific to *EN-Anxiety* (U=397, P= 0.41, one-tailed Wilcoxon Rank sum test, Fig. 6J). Thus, chronic social defeat stress increased *EN-Anxiety* activity in both susceptible and resilient groups of mice, which both exhibited anxiety in the EPM. Importantly, this stress-induced increase in *EN-Anxiety* was observed in the home cage context, which normally was considered a low-anxiety state.

We next quantified network activity in mice exposed to chronic mild unpredictable stress. In this paradigm, mice are repeatedly exposed to a series of stressors over eight weeks. Specifically, test mice are subjected to two stressors per day, one occurring during the light phase of their circadian rhythm cycle and the other during the dark phase. Stressors, including environmental stressors, food/water restriction, or physical restraint, were chosen according to a pseudo-randomized schedule. Exposure to this protocol induces altered reward and social behavior, as well as increased anxiety-related behavior in mice compared to their non-stressed controls ^45,46^. After verifying that chronic mild unpredictable stress induced open arm avoidance in the EPM (i.e., increased anxiety-related behavior; t_19_=2.37, P=0.018, one-tailed Welch’s t-test; see Fig. 6K), we quantified *EN-Anxiety* activity in stressed mice and non-stressed controls. We again hypothesized that stress exposure induces anxiety – and therefore higher *EN-Anxiety* activity – in contexts where anxiety is not observed under normal conditions. Thus, we quantified network activity from stress and non-stressed mice while they were in their home cage. Like chronic social defeat stress, exposure to chronic mild unpredictable stress increased *EN-Anxiety* activity (U=81, P=0.036, one-tailed Mann-Whitney U test, Fig. 6L). No such increases were observed in *EN-SocialAppetitive* in the stressed group, demonstrating that the stress-induced change in network activity was specific to *EN-Anxiety* (U=101, P= 0.60, one-tailed Wilcoxon Rank sum test, Fig. 6M). Together, these findings establish that two of the most widely utilized paradigms for modeling depression in mice converged on a common network-level signature that can be observed when mice are placed in a context normally associated with a low-anxiety state. These results also highlight the translational utility of *EN-Anxiety* for phenotyping mouse models of mood disorders.

## DISCUSSION

Preclinical models have played a role in the development of therapeutics for emotional disorders. These efforts would be greatly enhanced by the discovery of biological mechanisms that instantiate affective internal states in health and disease. Any such mechanisms must generalize across individual animals and affective contexts to achieve their true translational potential. Here, we employed multi-site electrical recordings in freely behaving mice, which were subjected to a collection of behavioral and experimental paradigms, all to discover and validate an electome network that encoded such a generalized anxious internal state. We reasoned that a putative anxious internal state could be observed at the intersection of many distinct paradigms used to model and induce anxiety in mice. Moreover, we believed that the unique features of these paradigms would enable us to disambiguate this anxious internal state from other internal states, such as arousal and reward, or other task-relevant variables (e.g., bright light, avoidance, etc., Fig. 7). Machine learning models trained solely using data from one anxiety paradigm failed to generalize to all three anxiogenic paradigms. On the other hand, a model trained using data from all three assays discovered a network reflecting a shared internal anxious state. Specifically, *EN-Anxiety* generalized to additional anxiety paradigms, including direct optogenetic interrogation of a circuit originating from a key network node (VHip) and a classic fear conditioning assay, highlighting its sensitivity. Finally, *EN-Anxiety* failed to encode multiple behavioral assays that induced rewarding and/or arousing (but not anxious) internal states, demonstrating its specificity. Thus, our multi-assay learning approach discovered a generalized anxious brain state.

**Figure 7:**
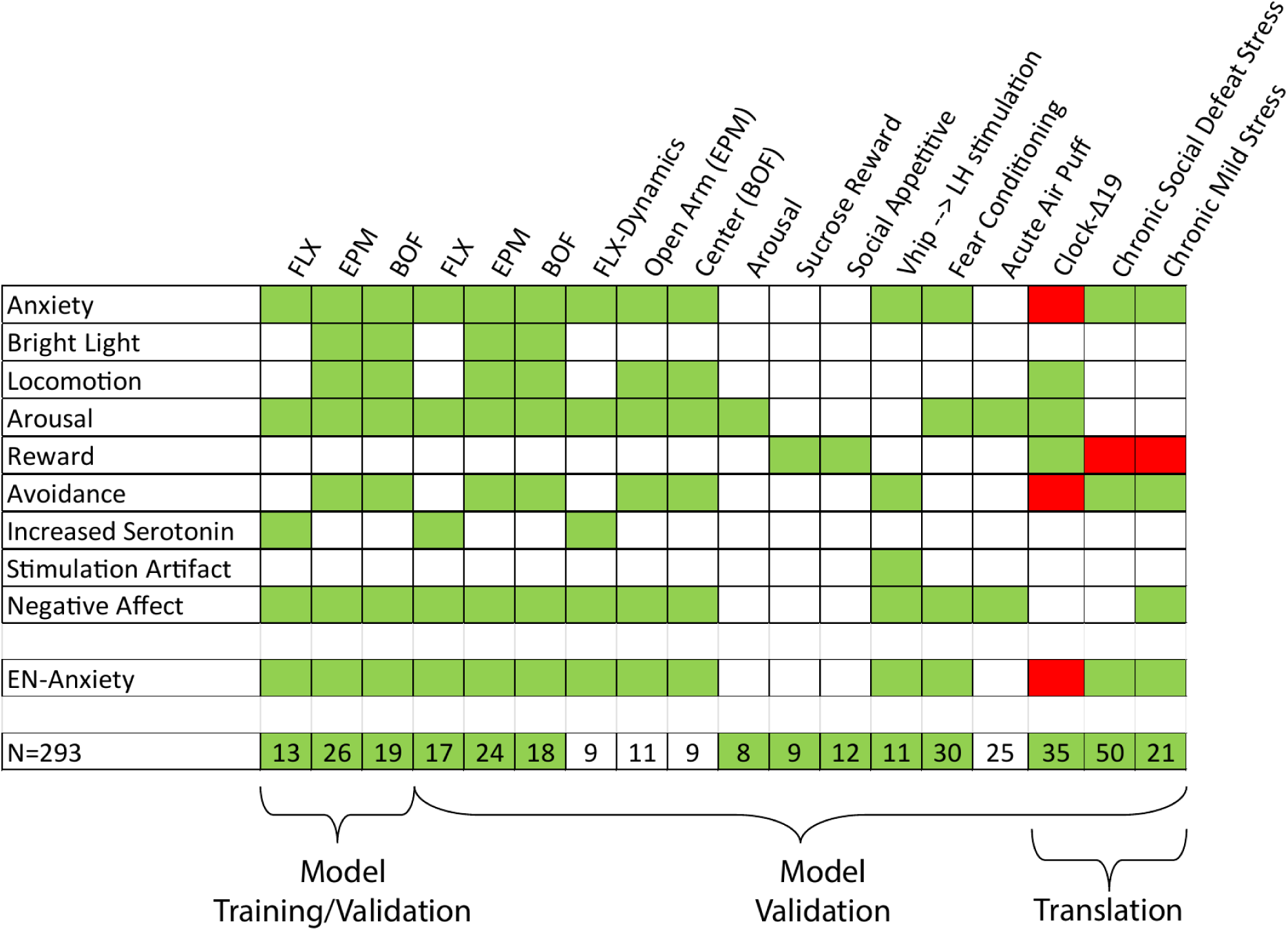
Conceptual framework utilized to discover and validate an electome network for anxious internal state. Affective/neurophysiological states and task variables are listed on the left of the table, and they are induced by behavioral and experimental manipulations (listed along the top). Manipulations that were hypothesized to increase the state/task variables are colored in green, while those hypothesized to decrease a state/task variable are colored red. Manipulations for which there is no clear prediction for the impact on the state are uncolored. Responses of *EN-Anxiety* to experimental conditions utilized throughout the study are shown in a separate row. Green and red boxes highlight conditions where network activity significantly increased or decreased, respectively. Numbers of mice used for each analysis are shown in the bottom row. New independent mice are highlighted in green.

In addition to robustly encoding multiple anxiety contexts, we also discovered that *EN-Anxiety* encoded location-dependent aspects of commonly utilized anxiety assays. Specifically, *EN-Anxiety* encoded occupancy of the avoidance zones of the EPM in the immediate past (Fig. 3F) and of the BOF in the immediate future (Fig. 3G). Thus, although these tasks are widely considered to be interchangeable assays for quantifying anxiety states in mice, we found that the pattern of *EN-Anxiety* encoding was distinct between the assays. Our behavioral analysis provided further support for the existence of key distinctions between the two anxiety-related tasks. Mice occupied the EPM avoidance zone throughout the assay (Fig. 3D), while mice showed greater occupancy of the BOF avoidance zone in the second half of assay vs. the first half (Fig. 3E). This latter observation was consistent with the high *EN-Anxiety* activity we observed during the first half of the BOF (Fig. 3B). Finally, we found no relationship between avoidance in both assays (Supplemental Fig. S2, bottom). Thus, while both assays engage key anxiety-related circuitry, the patterns whereby anxiety circuits shape behavior may be distinct across these two assays.

While each of our initial three paradigms could be encoded by LFP activity from at least one implanted region, no single brain region could independently encode an internal state shared across the three paradigms. Previous findings have established that a circuit involving Amy and VHip encodes anxiety in the EPM task in mice^31^. Indeed, we were also able to decode anxiety in the EPM and BOF solely using LFPs recorded from these regions (power in both regions, and the coherence and directionality measures between them). Yet, we were unable to capture an anxiety state shared across all three assays solely using LFP activity from this or any other pairs of brain regions recorded in this study. This suggests that activity within a given brain region or circuit can capture some behavioral/affective features of an individual assay, while failing to independently encode a generalized anxiety state shared across all three assays. For example, activity in a region/circuit may encode non-specific neural responses to the induction paradigms (e.g., sensing a bright light, or non-specific drug effects like large-scale serotonin increases in the brain), or behavioral features that correspond with anxiety in at least one of the three assays (e.g., locomotion). Because we employed different anxiety induction protocols (bright light, elevated arena, drug injection) and behavioral contexts (novel testing arena vs. home cage), we encouraged our machine learning strategy to discover a network-level, generalized anxiety state rather than circuits that encode specific features of each assay. Therefore, we assert that while many individual regions or circuits may contain assay-relevant information, the anxious brain state is optimally represented at the network level, where activity across many distinct brain regions/circuits is integrated at the sub-second timescale. Taken together, our findings highlight two important principles to help discover the neural architecture underlying affective states: 1) employ multiple distinct paradigms to discover the neural basis of generalized affective states, which helps to avoid finding a neural code that encodes features of only 1-2 specific assays; and 2) use neural activity acquired concurrently from multiple brain regions ^24^.

We do not contend that the learned *EN-Anxiety* provides a comprehensive description of the anxious internal state. Rather, we believe that this network-level state is also coupled with physiological changes across brain regions involved in sensory, motor, and homeostatic functions, and other circuits throughout the body (e.g., heart-brain and/or gut-brain interactions, etc.). It is also likely that several neural circuits outside of *EN-Anxiety* can converge to impact its activity. Indeed, we found that stimulating VHip projections in LH – a region that was not utilized to learn *EN-Anxiety* – increased *EN-Anxiety* activity. Thus, we assert *EN-Anxiety* provides a robust and objective neural measure of the internal state that mediates anxious behavior.

To test the translational utility of *EN-Anxiety*, we quantified network activity in a mouse model of mania and in two well-established preclinical animal models of depression based on chronic stress exposure. The *ClockΔ19* mouse model of mania exhibits predictive validity, as it shows increased reward drive and decreased anxiety-like behavior that responds to chronic lithium treatment ^38^. *ClockΔ19* mice exhibited normal *EN-Anxiety* network function (i.e., *EN-Anxiety* activity increased with exposure to the EPM), yet the network’s activity level in relation to these mice’s occupancy of the avoidance zone was blunted relative to wild type littermates. Thus, in addition to our prior work suggesting that the *ClockΔ19* mice’s deficits in anxiety behavior may be partly due to altered activity in reward circuits^15^, these findings argue that deficits in *EN-Anxiety* coding may play a role as well. Both chronic-stress models we tested exhibit predictive validity with depression, as they produce heightened anxiety-like behavior and an anhedonia phenotype that responds to chronic antidepressant administration ^43,47,48^. We observed increased *EN-Anxiety* activity in both depression models when mice were in a context that is otherwise characterized by low anxiety (i.e., their home cage), further arguing for their face validity as a model of stress pathology.

Taken together, our findings establish a brain electome network that encodes anxiety-related behavior in otherwise healthy animals and altered anxiety states in mouse models of mood disorders. Thus, we contend that *EN-Anxiety* presents a putative preclinical biomarker for phenotyping mouse models and for the development of anxiolytic therapeutics.

## MATERIALS AND METHODS

### Animal Care & Use

Male C57BL/6J (C57) mice were purchased from Jackson Labs at 6-8 weeks of age. *Clock*Δ19 mice were created by N-ethyl-N-nitrosourea mutagenesis that produced a dominant-negative CLOCK protein as previously described ^39,40^. After backcrossing >10 generations on a BALB/cJ background, *Clock*Δ19 mice and their wild type littermate controls were bred from heterozygous (*Clock*Δ19 -/*+*) breeding pairs. Male and female mice, 8-16 weeks old, were used for electrophysiological experiments presented in this study. Male CD1 mice (retired breeders) were purchased from Charles River Laboratories (Wilmington, Massachusetts) at 6 months. These mice were singly housed with enrichment except during their use in the chronic social defeat stress protocol. Unless otherwise specified, mice were housed 3-5 per cage, on a 12-hour light/dark cycle and maintained in a humidity- and temperature-controlled room with water available *ad libitum.* Most anxiety-related manipulations and behavioral tests were conducted with approved protocols from the Duke University Institution Animal Care and Use Committee, with the following two exceptions. The elevated plus maze (EPM) behavioral experiments in unimplanted *ClockΔ19* mice and their littermate controls were conducted at the University of Pittsburgh in compliance with approved protocols from the University of Pittsburgh’s Institution Animal Care and Use Committee. The EPM behavioral experiments in mice exposed to chronic social defeat stress were conducted at the University of Iowa in compliance with approved protocols from the University of Iowa’s Institution Animal Care and Use Committee. All experiments were conducted in 6-20-week-old mice, and in accordance with the NIH guidelines for the Care and Use of Laboratory Animals.

### Data Extraction and Processing

### Electrode Implantation Surgery

The electrode implantation surgery procedure has been described previously^49,50^. Mice were anesthetized with 1.5% isoflurane, placed in a stereotaxic device and metal ground screws were secured above anterior cranium (midline) and cerebellum (midline). A third screw was secured laterally, roughly half-way between the two other screws. 32 tungsten microwires were arranged in array bundles designed to target amygdala (Amy, 6 microwires), medial dorsal nucleus of thalamus (MD, 3 microwires), nucleus accumbens core and shell (NAc, 8 microwires), ventral tegmental area (VTA, 4 microwires), medial prefrontal cortex (mPFC, 8 microwires: 4 for cingulate cortex (Cg), 2 for prelimbic cortex (PrL), and 2 for infralimbic cortex (IL)), and ventral hippocampus (VHip, 3 microwires). They were centered based on stereotaxic coordinates measured from bregma (Amy: -1.4mm AP, 2.9 mm ML, -3.85 mm DV from dura; MD: -1.58mm AP, 0.3 mm ML, -2.88 mm DV from dura; VTA: -3.5mm AP, ±0.25 mm ML, -4.25 mm DV from dura; VHip: -3.3mm AP, 3.0mm ML, -3.75mm DV from dura; mPFC: 1.62mm AP, ±0.25mm ML, 2.25mm DV from dura; NAc: 1.3mm AP, 2.25mm ML, -4.1 mm DV from dura, implanted at an angle of 22.1°). The wires targeting PrL were built with a 0.25mm stagger anterior and a 0.5mm stagger ventral to the IL wires. Wires targeting Cg were built with a 0.6mm stagger anterior to the wires targeting PrL and a 1.1mm stagger anterior to the wires targeting IL. Animals were implanted bilaterally in mPFC and VTA. All other bundles were implanted in the left hemisphere (Supplemental Fig. S15). The NAc bundle included a 0.6mm DV stagger such that wires were distributed across NAc core and shell. We targeted basolateral amygdala (BLA) and central amygdala (CeA) by building a 0.5mm ML stagger and 0.3mm DV stagger into our Amy electrode bundle ^26^. Notably, these implantation sites have been homogenized across experimental preparations in the lab, enabling comparative analysis across previously and recently collected data sets. A metal ground wire was secured to the anterior and posterior screws, and the implanted electrodes were anchored to all three screws using dental acrylic. To mitigate pain and inflammation related to the procedure, all animals except those subjected to fear conditioning, chronic mild unpredictable stress, and chronic social defeat stress received carprofen (5mg/kg, s.c.). Injections were given once prior to surgery and then every 24 hours for three days following electrode implantation.

### Neural Electrophysiological Data Acquisition & Video Recording

Neurophysiological data were acquired using a Cerebus acquisition system (Blackrock Microsystems, Inc., Salt Lake City, UT). Animals were connected to the system using an M or Mu-32 channel headstage (Blackrock Microsystems, Inc., Salt Lake City, UT) and a motorized HDMI commutator (Doric Lenses, Quebec, Canada). Local field potentials (LFPs) were bandpass filtered at 0.5–250Hz and sampled/stored at 1kHz. All neurophysiological data were referenced to a ground wire connecting the ground screws above cerebellum and anterior cranium. Video recordings were acquired in real-time using NeuroMotive (Blackrock Microsystems, Inc., Salt Lake City, UT) and synchronized with neurophysiological data.

### Histological Confirmation

Histological analysis of implantation sites was performed using one of two protocols at the conclusion of experiments to confirm electrode placement. Animals were perfused with 4% paraformaldehyde (PFA, Electron Microscopy Sciences), and brains were harvested and stored for 24 hours in PFA. Brains were either processed on a cryostat or vibratome. 1) For cryostat: Brains were then cryoprotected with sucrose and frozen in OCT compound prior to being stored in -80°C. Brains were sliced at 35 µm using a cryostat and stained with either DAPI (AbCam) or cresyl violet (Sigma) using standard protocols. Slices were imaged at 4x and 10x magnification on a Nikon eclipse fluorescent microscope. 2) For vibratome: mice were perfused with 4% paraformaldehyde (PFA, Electron Microscopy Sciences) in PBS, and brains were harvested and post-fixed in 4% PFA and then transferred to PBS with 0.05mM sodium azide. Brains were sliced at 40 µm (Leica Vibrating Blade Microtome) and stained with Hoechst (Fishersci) containing mounting solution (9.6% Mowiol 4-88 (Sigma) in 24% glycerol, 0.M M Tris-Cl pH 8.5) on standard microscope slides. Slides were imaged at 4x and 10x with Olympus Slide Scanner (VS200).

### LFP Processing to Remove Signal Artifact

We employed a heuristic approach to eliminate recording segments containing non-physiological signals identically to previous works ^26,51^, and we paraphrase the processing procedure as follows: we first computed the signal envelope for each channel by utilizing the magnitude of the Hilbert transform. For any 1-second window in which the envelope surpasses a predetermined low threshold, we discard the entire segment if, at any point within that window, the envelope exceeds a second, higher threshold. The two thresholds were independently determined for each brain region. The high threshold was set at 5 times the median absolute deviation of the envelope value specific to that region. This choice was based on its approximate equivalence to 3 standard deviations from the mean in normally distributed data, while remaining robust to outliers. The low threshold was empirically established as 3.33% of the high threshold. If more than half of the window was removed for a given channel, we also removed the remaining portion of that window for that channel. Additionally, any windows where the standard deviation of the channel is less than 0.01 were excluded. This conservative procedure resulted in 10.7% of windows being removed.

### Feature Extraction

Feature extraction was performed identically to previous works ^26,51^, and we paraphrase the generation procedure as follows: LFPs were averaged across wires within the same region to generate a composite LFP measure. Signal processing was conducted using MATLAB (The MathWorks, Inc., Natick, MA). For LFP power, a sliding Fourier transform with a Hamming window was applied to the averaged LFP signal utilizing a 1-second window and a 1-second step. Frequencies ranging from 1–56Hz were analyzed. LFP cross-regional coherence was computed from pairs of averaged LFPs using magnitude-squared coherence, where coherence is a function of the power spectral densities of brain regions A and B and their cross-spectral densities.

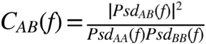

Spectral Granger causality features ^52^ were computed using the multivariate Granger causality (MVGC) MATLAB toolbox ^53^. The data underwent a high-pass Butterworth filter with a stopband at 1Hz and a passband at 4Hz. Directionality values for each window were calculated using a 20-order AR model through the GCCA_tsdata_to_smvgc function of the MVGC toolbox. Directionality values were determined for all integer frequency values within the specified range for all directed pairs of brain regions in the dataset^51^.

### Acute Fluoxetine Administration (FLX)

For the behavioral fluoxetine experiments, mice were pseudorandomly assigned to receiving an injection of either fluoxetine or saline 30 minutes prior to being placed on the elevated plus maze (EPM). Fluoxetine (Sigma) was made up in 0.9% NaCl to a concentration of 1mg/mL and then injected at 10mL/kg for a final concentration of 10 mg/kg, i.p.^28^. Physiologic saline injection was injected at 10 mL/kg as a control for injection volume. Animals were habituated to i.p. injections daily for 1 week prior to behavioral testing.

For electrophysiological recordings, animals used for training the final multi-assay model followed a standard pharmacological crossover design. Though fluoxetine only has an 8-hr half-life in mice, a lengthy washout period of two weeks was used to ensure no traces of the drug remained. Specifically, after habituation to the experimental room for 1 hour, mice were pseudorandomly assigned to receive a fluoxetine or saline injection. Neural recordings were then obtained for an hour. Two weeks later, animals underwent a second 1-hour recording session after receiving the other treatment. To test the final model, we utilized a protocol in which two recordings were performed at a much closer interval. Specifically, after habituation to the experimental room for 1 hour, nine new mice were treated with saline and neural data was recorded for an hour. Four hours later, mice were subjected to a second recording session immediately following treatment with fluoxetine.

### Elevated Plus Maze (EPM)

The EPM assay is widely employed to measure anxiety behavior in mice^54^. The following procedure was utilized for all studies except for the experiments conducted in unimplanted *ClockΔ19* mice and their wild type littermate controls. The EPM is comprised of four arms arranged in a cross shape, each measuring 30.5cm in length and 5cm in width, positioned at a height of 91.4cm from the floor. Additionally, there is a central region measuring 5cm by 5cm. Among the arms, two are designated as ‘closed,’ enclosed by walls that are 16.5cm in height on three sides, while the other two are ‘open’ and surrounded by a low piece of tape, approximately 1mm in height.

Two days prior to testing, mice were gently handled in the experimental room for roughly 1 minute per animal. Following gentle handling, mice were habituated to the room for 1 hour in a testing ‘home cage’. Afterwards, mice were returned to group housing in their original home cage. This procedure was repeated one day prior to experimental testing. On the testing day, mice were habituated to the experimental room in their individual testing home cage for 1 hour. Mice were then connected to the recording system and habituated for an additional 10 minutes. Following 5 minutes of neural and video recordings from an overhead camera, mice were placed in the center of the EPM facing one of the closed arms. Neural recordings and video data were acquired for an additional 5-10 minutes. Testing was performed at 175 lux during the light cycle. A subset of the mice (N=15) utilized to train and test the multi-assay model were previously used in a prior study to identify an electome network that encoded social appetitive behavior^26^. In that study, mice were recorded during social exploration, and activity in the discovered network was also evaluated as mice explored an EPM.

For experiments conducted in unimplanted *ClockΔ19* mice and their wild type littermate controls, adult male and female *ClockΔ19* and wild type mice (8-14 weeks; N=17 per genotype, balanced across sex) were habituated to the testing room for 1 hour prior to testing. Chambers were cleaned with 70% ethanol and allowed to dry between animals. The elevated plus maze (EPM) consisted of two plastic open arms perpendicular to two closed arms (30 × 5 cm) and was elevated above the ground (60 cm). Mice were placed in the center of an EPM, and behavior was recorded for 10 minutes. Ethovision video tracking software (Ethovision, Noldus, Leesburg, VA, USA) was used to quantify the time spent in the open and closed arms, the number of entries into the arms, and the distance travelled. Manual scoring was also conducted for verification. Testing was performed at 20 lux.

### Bright Open Field (BOF)

The BOF assay is also widely employed to measure anxiety behavior in mice^54^. This assay consists of a square arena (46 cm x 46 cm x 30 cm) in which the innermost third (i.e., ‘center zone’) is considered more anxiogenic zone than the outermost two thirds (i.e., ‘periphery zone’). Mice were habituated to the experimental room in an individual testing home cage using the same procedure described for the EPM. On testing day, mice were connected to the recording system and 5 minutes of neural and video data (from an overhead camera) were acquired while mice were in their individual testing home cage. Mice were then placed in the periphery of the BOF, and an additional 5 minutes of neural and video data were acquired while mice freely explored the arena. Testing was performed at 125 lux, during the light cycle.

### Heart Rate Monitoring

Unimplanted mice (N=7-9 male and 7-8 female mice per assay, i.e., FLX, EPM, BOF) were anesthetized using 1% isoflurane, and the fur around their neck region was removed using potassium thioglycolate (Nair Body Cream, Church & Dwight Co. Ewing, NJ). Awake mice were then habituated to the monitor cuff using a small or extra-small CollarClip (Starr Life Sciences Corp, Oakmont, PA) for 1 hour/day for 7 days in their home cage environment. Additionally, the mice used for the FLX assay experiment were habituated to daily i.p. injections using saline.

On testing day, the CollarClip sensor was placed around the neck and mice were allowed to acclimate for 45 minutes. Measurements were then collected using the MouseOx Plus (Starr Life Sciences Corp, Oakmont, PA). For EPM and BOF, baseline data was collected for 5 minutes while mice were in their home cage, and for an additional 5 minutes as mice were subjected to the EPM or BOF. For the FLX assay injections, mice were first injected with saline and recorded for 1 hour. After at least 5 hours, mice were injected again with fluoxetine (10 mg/kg, i.p.) and recorded for an additional hour. Data points less than 250bpm were attributed to movement artifact and removed from the data. Data was acquired at 1Hz, and mean heart rate values were calculated for the low- and high-anxiety contexts for each assay. A one-tailed t-test was used to evaluate whether heart rate was higher in the high- vs. the low anxiety context.

### Sleep State Identification

We utilized data collected as part of a previous study^33^. In brief, eight mice were implanted with the same brain targets as described above as well as a wire placed in the trapezius muscle to acquire electromyography (EMG) data. Mice were singly housed, habituated to the recording room for 2 days, and neural data was then collected for 24 hours. EMG power (root mean squared of the spectral amplitude of 30–56Hz) and a ratio of prelimbic cortex activity (2–4.5 Hz/2–9 Hz) were used to classify 3 sleep-wake states based our previously published clustering approach in mice^55^. High EMG activity but low ratio of prelimbic cortex activity corresponded to the awake state, low EMG activity but high ratio prelimbic cortex activity corresponded to NREM sleep, and low EMG activity and low ratio of prelimbic cortex activity corresponded to REM sleep. Data that was not classified within one of these three states was classified as ‘unlabeled’ (Fig. 4A).

Using this classified data, we selected large time segments for which neural data was classified as wakefulness or REM sleep. Specifically, awake intervals were defined by at least 30 seconds of awake-labeled data with at least 5 minutes of >95% awake labels on both sides of the data segment (Fig. 4B, top). REM sleep intervals were defined by at least 60 consecutive seconds that were labeled as REM sleep (Fig. 4B, bottom). LFP data from these intervals were then projected into *EN-Anxiety* and *EN-SocialAppetitive*^26^. All animals were recorded from each of the brain regions that composed both networks.

### Delayed sucrose reward apparatus, training, and task

The delayed sucrose reward task was modeled after a prior test in which mice had to remain in a spatial location to receive a food reward ^56,57^.

The task chamber was constructed from Lego Duplo® pieces of varied color, shape, and size. The apparatus had approximate dimensions of 48cm wide x 35cm deep x 30cm tall, and each wall was visually distinct. A nose poke detector was in the center of each wall, placed 1cm above the floor, with an LED light directly above each nose poke detector. Four fluid dispensers were calibrated to release 5µL of 10% sucrose directly into each nose poke detector. The reward for three of the ports were also flavored with pumpkin, almond, or orange oil. The location and reward type remained fixed throughout each phase of experimental testing for all animals. During the task, the chamber was illuminated to 30 lux. The system was also equipped with speakers and an audiometer, and reward cues were played at 68dB. Signals from the nose poke detectors, LED lights, fluid dispensers, and audiometer were digitized and stored in parallel with our neural recordings.

After 7-14 days of recovery from surgical implantation of electrodes targeting the same regions composing *EN-Anxiety*, C57 male mice (N=9) were food-deprived to 90% of their free-feeding body weight. During a training session, a mouse was connected to a recording cable and placed in the task apparatus. The training procedure is as follows:

- Stage 1: On day 1, mice freely accessed the testing chamber for 60 minutes. Each poke into a nose poke detector triggered a 500ms tone at 4000Hz and 5µL of reward release directly into the poke detector. This stage was repeated over 2 days.
- Stage 2: On days 3-4, mice were placed into the recording chamber together with their cage mates, without a recording cable. Mice were then allowed to freely explore the recording chamber for 120 minutes.
- Stage 3: On day 5, mice resumed individual training, during which they advanced in task difficulty after meeting specific criteria.
- 3a: Each detected poke activated a 500ms, 4000Hz tone and released a 5µL reward at the beginning of the tone (onset).
- 3b: Each detected poke activated a 500ms, 4000Hz tone and released a 5µL reward at the end of the tone (offset).
- 3c: Each detected poke activated a 500ms, 4000Hz tone and released a 5µL reward at tone offset if a mouse remained in the nose poke detector.
- 3d: Each detected poke activated a 500ms, 4000Hz tone and released a 5µL reward 1 second after tone onset if a mouse remained in the nose poke detector.
- 3e: Each detected poke activated a 500ms, 4000+387Hz tone, and a second 500ms 4000Hz tone, 1 second after onset of the first tone. A 5µL reward was released 1.5 seconds after onset of the first tone if a mouse remained in the nose poke detector.
- 3f: Each detected poke activated a 500ms, 4000+387Hz tone, and a second 500ms 4000Hz tone, 1 second after onset of the first tone. A 5µL reward was released at 2 seconds after onset of the first tone if a mouse remained in the nose poke detector.
- 3g: Each detected poke activated a 500ms, 4000+387+387Hz tone, and a second 500ms 4000+387Hz tone 1 second after onset of the first tone, and a final 500ms, 4000Hz tone 1 second after onset of the previous tone. A 5µL reward was released 2.5 seconds after onset of the first tone if a mouse remained in the nose poke detector.
- This training pattern continued until mice passed training with a 5-second delay. For these trials, successive tones of decreasing frequency were played, one at the beginning of each second, and mice received reward if the poke hole was activated for the entire test interval.
- A mouse passed a training stage when it completed 120 rewarded pokes in one day, or 120 rewarded pokes in two consecutive days and the second day reward count was greater than or equal to the reward count of the first day. Mice regressed to a prior training stage if they failed to complete a stage after five days, or if they received fewer than twenty rewards during a session. The data utilized for our electrophysiological analysis was acquired from 9 C57 male mice after they completed training at the 5-second delay.

### Social Preference Assay

A previously published data set^26^ was used to assess the impact of social interaction on *EN-Anxiety* activity here. Briefly, mice implanted with electrodes at the same brain coordinates utilized for this study (N=12 C57 mice) were allowed to explore a rectangular arena (61cm × 42.5cm × 22cm, L×W×H) for 10 minutes. Two clear plexiglass walls divided the area into two equal chambers. Each chamber contained a circular holding cage (8.3cm diameter and 12cm tall) containing either a novel object or a C3H target mouse matched for sex and age. Data was collected across 6-10 testing sessions/mouse. Video data was tracked using Bonsai Visual Reactive Programming software, and network activity was analyzed for periods in which mice were within ∼5cm of the novel object or target mouse. This neural data was projected into *EN-Anxiety* (see "Projecting Data into a Previously Learned Network"). Six of the 12 experimental mice were subjected to the EPM assay after the social preference assay, and their EPM data were included as part of the training/validation/testing data for the multi-assay network model.

### Optogenetic Stimulation and Behavioral Testing

Mice were injected with AAV5-ChR2-eYFP into VHip at 7 weeks old. Eight weeks after the injection, mice were implanted with a fiberoptic cannula either targeting right lateral hypothalamus (LH: −1.95AP, 0.5ML, −4.75DV; N=13 C57 mice) or right basal amygdala (Amy: -1.7AP, 3.0ML, -4DV; N=18 C57 mice). Behavioral experiments (BOF, real-time place preference) were conducted after 2 weeks of recovery; experimental details are listed below.

#### Bright Open Field (BOF) testing

Mice were habituated to the experimental room and fiberoptic tethering for 2 days prior to testing. The BOF arena was used for testing, where the innermost third was designated as the center zone. The center area was illuminated to 600 lux. On testing day, mice were connected to the fiberoptic patch cord and immediately placed into the open field. All mice were stimulated with yellow light (593.5nm, OEM Laser Systems, Model No. MGL-F-593.5/80mW, 20Hz, 5ms pulse, either 1mW for Amy or 5mW for LH) while they explored the BOF chamber for three minutes. Mice were then pseudorandomized to receive either blue (473nm, CrystaLaser, CL473-025-O, 5ms pulse, either 1mW for Amy or 5mW for LH) or yellow light and stimulated for the next three minutes. The location of mice was tracked using Ethovision XT17 (Noldus). Behavior during this second stimulation interval was analyzed using a one-tailed t-test to test whether greater avoidance was observed during blue light stimulation.

#### Real-Time Place Preference

One week after BOF testing, mice were pseudorandomly re-assigned to either a blue or yellow light group. A two-chamber arena (60cm x 42cm x 22cm) was used where one side was pseudorandomly assigned to be the stimulation side for each mouse. Mice were connected to the fiberoptic cable and placed into the arena for 20 minutes. Real-time tracking (Neuromotive, Blackrock Neurotech) was used to activate the lasers (same parameters as BOF testing) when a mouse was detected on the stimulation side. Movement was tracked using Ethovision XT17 (Noldus), and % time on the stimulation side was analyzed using a one-tailed t-test to test whether greater avoidance was observed during blue light stimulation.

### Optogenetic Stimulation and Electrical Recordings

We modeled previously published methods for targeting the VHip → LH circuit ^20^. Specifically, mice were anesthetized with 1.5% isoflurane and placed in a stereotaxic device. A 33-gauge Hamilton syringe was used to infuse 0.5 μl of AAV5-ChR2-EYFP at a rate of 0.1 μl/min into right VHip (-3.16mm AP, 3.3mm ML, -3.75mm DV from dura; N=11 mice). Eight weeks later, mice were implanted with recording electrodes using the procedure and brain targets described above (*"Electrode Implantation Surgery"*). These electrodes included a bundle that was used to target the right lateral hypothalamus (LH: −1.95AP, 0.5ML, −4.75DV). A 100µm diameter fiberoptic (Doric Lenses) fiberoptic cannula was built into the LH bundle with the tip situated 250µm above the tip of the LH microwires bundle ^58,59^. *In vivo* recordings were conducted after 2 weeks of recovery. Mice were habituated to the experimental room/setup for the two days preceding experiments. As a control experiment, the same procedure was used to test the impact of VHip → amygdala stimulation, with the exception that the optrode was implanted to target right amygdala (Amy: -1.7AP, 3.0ML, -4DV; N=9 mice) instead of right LH. We also utilized a light intensity of 1mW because our initial experiments utilizing a light intensity of 5mW induced electroencephalographic and myoclonic seizures.

Mice were connected to the recording system using a fiberoptic patch cable and a 32-channel M headstage and then placed in a new ‘testing’ home cage for 1 hour. On testing day, mice were connected and placed in the same testing home cage. After 40 minutes of additional habituation, neural data was recorded for 20 minutes. Mice were then stimulated with blue or yellow light for 10 minutes. Light stimulation was delivered with 5ms pulses at 20Hz, 5mW, and verified using a power meter (Thorlabs, PM100D). Mice were pseudorandomized to stimulation with either blue (473nm, CrystaLaser, CL473-025-O) or yellow light (593.5nm, OEM Laser Systems, Model No. MGL-F-593.5/80mW).

One week later, mice were subjected to a second recording with the other laser, using the same protocol described above. Thus, each mouse was stimulated with blue and yellow in pseudorandomized order across the two sessions.

### Fear Conditioning

Mice were implanted with electrodes as described above (*"Electrode Implantation Surgery"*; N=36 male C57 mice). Following a two-week recovery period, mice were trained in a classic cued fear conditioning paradigm during which an auditory tone (conditioned stimulus; CS) was paired with an aversive air puff (unconditioned stimulus, US). The CS consisted of a 30-second, 10 kHz, 80dB, continuous auditory tone that was generated using MATLAB. The US consisted of a 2-second, 40 PSI, air puff that was introduced through 4 pumps, one built into each of the testing chamber’s walls. Mice were pseudorandomly assigned to 2 groups: conditioned (CS+; N=17) and control (CS-; N=19). Behavioral testing was conducted in two different behavioral contexts (context A and B). Context A was a 10"×10"×11" (L×W×H), striped chamber made of alternating black and white Legos®. Context B was a 6"×12"×11" chamber, with walls consisting of mixed colored Legos. Context A had a smooth floor, while context B had a textured floor.

Prior to conditioning (Day 0), mice were habituated to the behavioral room for 2 hours. On Day 1, mice were connected to the recording system and placed into Context A for 2 minutes. The fear-conditioned group was exposed to 7 trials of the CS. The US was presented during the last 2 seconds of each tone, and there was a pseudorandom interstimulus interval ranging from 60-120 seconds. The control group was exposed to 7 trials of the CS without the US. Each group remained in Context A for 1 minute after the last trial concluded. Neural and video data were collected throughout the trials. We also collected a continuous signal corresponding to the onset and offset of the CS.

Mice were then exposed to a cued recall session on Day 3. Here, mice were connected to the recording system and placed into Context B. After 3 minutes, mice were presented with the CS tone continuously for 3 minutes. Neural and video data was recorded throughout this interval, and the freezing behavior was quantified using Ethovision X12 (Noldus, Wageningen, the Netherlands) to detect the percentage of time during the CS presentation that the animal did not move. A subset of the conditioned mice (N=5) was also exposed to an extinction protocol on Day 2, where mice were presented with the CS, but not the US, in Context A. Since our objective was to quantify neural responses to fear conditioning on Day 1, and exposure to the one-day extinction protocol had no impact on freezing behavior on Day 3 (t26=0.11, P=0.92, two-tailed unpaired t-test), we pooled this subset into the CS+ group for the analyses presented in the text. All implanted mice were used for behavioral analysis. Three CS+ mice and two CS-mice were removed from neurophysiological analysis due to poor signal quality.

### Electrophysiological analysis of *ClockΔ19* mice

*ClockΔ19* mice and their wild type littermate controls were implanted and recorded in the EPM assay as described above (22-23 mice/genotype, balanced across sex). Six mice had poor electrophysiological signal prior to testing and were thus completely removed from the study. Four mice had poor quality video recordings and were thus removed from behavioral analysis (i.e., 17-18 per genotype, balanced across sex, were used for behavioral analysis). Two of these mice had poor electrode placements and were thus removed from subsequent video electrophysiological analysis. Thus, a total of 33 mice (16-17 per genotype) were used for EPM location-based electrophysiological analyses. Since the start time of the EPM task was noted for each experiment, the four mice with poor video quality were included in the analysis of network activity induced by task exposure (home cage vs. EPM across genotype).

For the EPM location analysis, the center of mass for each mouse was identified using DeepLabCut^60^. This center of mass was then used to label the location of 1-second data windows as the center, open arm, or closed arm based on which location they spent more than half of the second. These center of mass-based labels were utilized for the analysis of closed arm/safe zone network activity in EPM. We also scored the location of the recording headcaps of mice using Ethovision. Because of the reduced size of the center location, we used the 1-second data windows for which both the center of mass and the headcap-based locations were labeled as the center of the maze. This approach ensured high specificity for the windows labeled as the center.

### Chronic Social Defeat Stress (cSDS)

These methods parallel those described in our prior work ^27,59^. Data for our electrophysiological and behavioral analyses were obtained from two different cohorts of implanted mice. Behavioral data on the EPM was assessed from mice implanted in a different set of brain regions than those used for this study. Neurophysiological recordings in the home cage were obtained from mice implanted in the same brain regions utilized in this study. Data from a subset of this latter group of mice were previously used in our published work ^26^.

Six- to seven-week old male C57 mice (N=50) were implanted with electrodes as described above (*"Electrode Implantation Surgery"*). Stress experiments were initiated two weeks after surgical recovery. All mice were singly housed prior to being subjected to chronic social defeat stress (cSDS). We modeled our protocol after previously published work ^43,47^. Implanted animals were pseudorandomly assigned to control (N=16) or stress groups (N=34), such that cagemates were distributed across groups. Singly housed male retired-breeder CD1 (Charles River) mice were used as resident aggressors for 10 days of social defeat. Briefly, mice in the stress group were exposed to a different CD1 aggressor for 5 mins each day and only removed early in the event of serious physical injury, which never happened for more than two stressed mice per 10 days of social defeat. Defeats were run in dim light conditions (∼40-50 Lux). After 5 mins, C57 mice and CD1 aggressors were separated with a perforated divider for 24 hours. Control C57 mice were housed with another C57 mouse in a similar cage apparatus. Their C57 cage mates were rotated each day. This process was repeated for a total of 10 days. Triage was performed on stressed animals following each day of defeat to check for and treat any wounds. After this check, the lights were turned off. Mice that exhibited significant injuries during social defeat stress were removed from the study. Neural recordings were obtained from control and stressed mice in their home cage one day following completion of the cSDS protocol. Mice were subjected to these recordings in their home cage, immediately followed by recordings during a forced interaction test. This assay has been previously described^27,59,61^, and only data from the home cage recordings are utilized in the present study.

A choice interaction test was used to categorize stressed mice as susceptible or resilient ^42,43,59,61^. This assay was performed 2 days after the last social defeat session, during the dark cycle of each mouse. Testing was conducted in a room with two red lamps facing the ceiling (2-10 lux). All animals were habituated to the room for approximately two hours prior to the start of testing. Experiments were pseudorandomized and balanced to include alternating control and stressed mice spread throughout the duration of experiment. For each trial, the experimental mouse was placed in the center of an opaque white 18" x 18" box, with 18" high walls and a wire-mesh sub-chamber at the center of one wall for 150 seconds. Then a CD1 mouse (low/non-aggressive) was placed in the enclosed sub-chamber, and the C57 mouse was placed back in the box for 2 minutes and 30 seconds. Behavior was recorded for the entirety of each trial. Stressed mice that showed higher interaction time with the empty sub-chamber than the sub-chamber containing the CD1 mouse were defined as susceptible. Mice that showed higher interaction time with the sub-chamber containing the CD1 were defined as resilient. Between mice, all chambers were cleaned with Super Sani-Cloth germicidal disposable wipes (PDI, Orangeburg, NY) or 70% ethanol and dried with kimwipes. Data was analyzed using Noldus Ethovision version 15.

Mice subjected to cSDS and their non-stressed controls were tested in the EPM 12 days following the last social defeat session. Animals were tested during their dark cycle in a dark room with two red light lamps facing the ceiling (2-10 lux at the surface of the behavioral arena). Animals were acclimated to the room for >1 hour prior to starting experiments. For each trial, animals were placed in the center of the apparatus facing the same side and allowed to explore the maze for 5 minutes. Afterwards, the animal was removed from the apparatus and placed back into its home cage. The trials were pseudorandomized and balanced with alternation of control and stressed animals. After each session, the EPM was thoroughly cleaned with Super Sani-Cloth germicidal disposable wipes (PDI, Orangeburg, NY) and dried with kimwipes. Data was analyzed using Noldus Ethovision version 15.

### Chronic Mild Unpredictable Stress

We modeled our chronic mild unpredictable stress protocol after previously published work ^45,46^. C57 male mice (N=21) were implanted at age 7-9 weeks with electrodes to target the same brain regions utilized to learn the anxiety-related networks. Two weeks later, cages of mice were pseudorandomized into a stress or control group. Control mice were subjected to gentle handling twice a week. The stress group was exposed to 2 aversive experiences each day – one during the light cycle and one during the dark cycle – for eight weeks, as previously described ^45^. The stressors were as follows:

- physical restraint – mice were placed in a ∼50mL plastic cone (with openings for breathing on both ends) for 1 hr
- shaking – a cage of mice was placed on an orbital shaker for 1 hr at 60 rpm
- overnight illumination – mice were exposed to regular room light during the 12-hr dark cycle
- inverted light cycle – mice were exposed to dark-cycle room conditions during the light cycle and light conditions during the dark cycle
- tilted cage – cages were tilted at a 45° angle for 12 hrs
- strobe – mice were placed in a room with a strobe light during the dark cycle for 12 hrs
- wet bedding – cage bedding was saturated with water for 12 hrs
- soiled rat bedding – cage bedding was replaced with used rat cage bedding for 3 hrs
- cold exposure – mice were placed in a cold room (4°C) for 1 hr
- missing bedding –bedding was completely removed from the cage for 12 hrs
- food and water restriction – food and water were removed for 12 hrs during the dark cycle
- overcrowding – cage space was reduced by 50% for 12 hrs during the dark cycle

> Stressors were presented in pseudorandomized order, solely ensuring > that animals did not receive the same stressor two days in a row. Body > weight was monitored once a week to ensure mice didn’t lose more than > 10% body weight during the stress proposal.

### Model Selection and Training

#### Label Assignment for Training Datasets

To make use of supervised machine learning methods, per-sample anxiety state labels must be assigned for our training assays. For the acute FLX assay, we accounted for drug activation time and assigned all time points within the last 30 minutes of the 1-hour recording to either a high- or low-anxiety state after mice received fluoxetine or saline, respectively. For the EPM and BOF assays, anxiety states within the assay can be ambiguous. To prevent mislabeling of anxiety states in these assays, we assigned all time points for which mice are in the EPM or BOF as a high-anxiety state. We then labeled recordings taken while the mice were in their home cage environment as a low-anxiety state. With this formulation, all three training assays had the same labeling nomenclature of high- and low-anxiety states regardless of the anxiogenic assay, allowing for easy combination and comparison during model training.

#### Training and Test Splits

Once features were extracted for the FLX, EPM, and BOF assays, the complete data set from an individual mouse was placed into one of two groups of data: training data and test data. These splits were performed per mouse such that all its data were contained in the same group, including if an individual mouse had data from multiple assays. Splitting by mouse is critical as it prevents a machine learning model from simply learning the identity of a mouse in the training data to achieve inflated performance on holdout data^62^. The training data were used for model development, where many sets of hyperparameters and model formulation were tested through a cross-validation procedure. Test data were kept as true holdout data, which we did not observe or test our model on until the final model architecture was determined. Once final model parameters were determined through cross-validation, the full training data set was used for a final training run, and the output was then tested on the true test (holdout) data. Training and test groups had: 13 and 9 male mice respectively for the FLX assay; 26 and 11 male mice respectively for the EPM assay; and 19 and 9 male mice respectively for the BOF assay. 17 of these training mice and 7 test mice were shared across 2 assays (EPM and BOF). There was a single male test mouse that was exposed to all three assays. We also performed independent testing in female animals. Test groups included 13 female mice for the EPM assay, 9 female mice for the BOF assay, and 8 female mice for the FLX assay. 3 female mice were shared between the FLX and BOF assays (mice were tested in the BOF assay first), and 3 female mice were exposed to all three assays. All mice used in multiple assays were exposed to FLX last.

#### Discriminative Cross-Spectral Factor Analysis (dCSFA-NMF)

Discriminative Cross-Spectral Factor Analysis - Nonnegative Matrix Factorization (dCSFA-NMF) is a machine learning framework for discovering key predictive factors relevant to a behavioral assay or emotional state of interest ^25^. This method has been used previously to detect brain networks corresponding to stress and social activity in mice using LFP data^26^. Similar to other factor models that have been used in neuroscience, such as principal component analysis (PCA), independent component analysis (ICA), and nonnegative matrix factorization (NMF), dCSFA-NMF identifies underlying components that are interpreted to be networks of connectivity. The superposition of these networks then explains the observed neural activity. While the previously mentioned unsupervised methods can identify networks of neural activity, it is unlikely that anxiety and other emotional states are directly related to one of these dominant networks. Thus, dCSFA-NMF makes use of a supervision component to ensure that one or more of the learned networks are correlated with a behavioral measure or emotional state of interest, meaning that it learns networks that both explain the observed neural activity and decode conditions of interest.

Rigorously, the model learns *K* fixed components *W* ∈ *ℝ*^*K* × *M*^ that can reconstruct observed data *X* ∈ *ℝ*^*N* × *M*^ using an array of network activity scores *s* ∈ *ℝ*^*N* × *K*^ such that *X* = *sW*. The observed data *X* are formed from the features extracted from the LFP signals (see *"Feature Extraction"* above), where each window of data *x_n_* ∈ *ℝ*^*M*^ represents the concatenation of power, coherence, and directionality features. *W* and *s* are constrained to be positive since the features of use – power, coherence, and directionality – observed in *X* are also nonnegative. Network activity scores are inferred from the observed data using an encoder function *s* = *f_θ_*(*X*), which can take the form of a neural network or linear model. The activity scores *s_s_* ∈ *ℝ*^*N* × *Q*^ of the *K* ≥ *Q* ≥ 1 supervised components are then used in a logistic regression model *f_ϕ_* to predict the behavior of interest *y* = *f_ϕ_*(*S_s_*). We constrain our predictions to use a sparse combination of all networks, namely only the supervised networks, to narrow the scope of our network discovery and reduce the total number of comparisons. The parameters of the model are then optimized using the loss function,

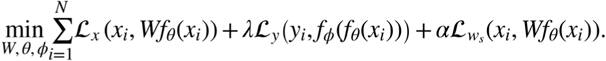

Here, *ℒ_x_* is the reconstruction error between the original power, coherence, and directionality features and those features generated by the product of our network scores and networks, *sW*. In this work, we make use of the Mean-Squared-Error (MSE) function. Our predictive loss *ℒ_y_* is a binary cross-entropy loss and penalizes our model for incorrectly predicting the behavioral state of each window. The impact of the predictive loss can be tuned using the hyperparameter *λ*. Lastly, we impose a second reconstruction loss, *ℒ_w_s__*, on the supervised network scores. This reconstruction loss prevents our neural network encoder *f_θ_* from learning an uninterpretable near-zero noise embedding for the supervised scores, which predicts the condition of interest (i.e., which condition of an anxiety assay a mouse is in) well with little to no contribution to explaining neural activity. This loss can be formulated as another MSE loss between the outer product of the supervised network activations and the supervised electome network and the features. Alternatively, this loss can be formulated as a penalty to drive the supervised network scores to reconstruct the residual of the unsupervised networks and features^63^. We used the latter in our analysis.

#### Performance Metrics -- Predictive Modeling

To evaluate the predictive performance of our model, we used the receiver operating characteristic curve - area under the curve (ROC-AUC). This metric is common in machine learning literature and is closely related to the concordance index for evaluating discrimination between binary labels. In our setting, the binary labels correspond to periods classified as a high-versus a low-anxiety state (e.g., EPM vs. home cage in the behavioral assays). AUC is a rank-based metric that quantifies the probability that a randomly chosen high-anxiety period receives a higher summary value than a randomly chosen low-anxiety period. AUC has values in the range [0,1] where AUC=0.5 indicates chance-level discrimination. AUC=1 perfect separation of the two binary states, and AUC=0 indicates perfect separation with the label ordering reversed. That is, the summary values used in the metric are consistently ordered such that low-anxiety periods receive higher values than high-anxiety periods.

For evaluating our models, we obtained an AUC for each mouse (e.g., how well does the model discriminate between a high-versus a low-anxiety state) and then reported the data group mean and standard error of the mean for each assay (e.g., FLX, EPM, BOF). Reporting AUC for each mouse rather than one single AUC value over all windows is more reflective of discriminative ability due to the differing base anxiety levels between mice. Additionally, many of our behavioral contexts have varying neural recording lengths; by reporting AUC values for each mouse separately, we addressed the possibility of our model overfitting to the neural activity of mice in paradigms with longer recordings and therefore more samples.

When we evaluate the AUCs reported over mice, we use a *t*-test to compare whether the mean AUC is different than the null value of 0.5. If a single network or prediction is evaluated, significance is evaluated by a Mann-Whitney-U statistical test compared to the chance-level discrimination of 0.5.

#### Performance Metrics -- Generative Modeling

We evaluated how well our models explained neural activity by quantifying the mean-squared-error (MSE) of the model’s predicted power, coherence, and directionality features and the originally observed values.

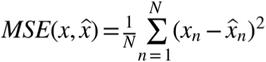

During model training, we weighted the reconstruction of each of our feature types (power, coherence, directionality) by their prevalence, such that power holds equal importance to coherence and directionality despite representing a smaller number of features.

#### Performance Metrics -- Model Consistency

In addition to our ability to reconstruct neural data and discriminate anxiety levels, we want our networks to be reliable and consistent, in the sense that they would be robust to minor perturbations of the data. Therefore, we measured how similar our electome networks of interest (the learned values of *W* corresponding that are predictive) are when learned from slightly different subsets of training data, which we refer to as "representation consistency"^63^.

To evaluate representation consistency in our model, we created a representation consistency distance score based on the weighted average cosine distance formula between vector representations of electome networks. Cosine distance calculates the angular separation between two vectors and takes values in [0, 1] for nonnegative vectors, where 0 represents perfect alignment and 1 represents orthogonal vectors. The cosine distance between two vectors, *A* and *B* is given by:

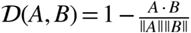

During the training process, we independently learned model parameters on two subsets of data (described below). After learning, we matched networks between each set of data to determine how similar the network in the first subset of data is to its uniquely matched network in the second subset of data (details given in the next paragraph). Instead of averaging directly over resultant cosine distances, we chose to weight each unique pairing in the average by the impact that a network score had on the prediction relative to the other networks.

In other words, a network that does much of the prediction on whether an animal is in a high- or low-anxiety state will be weighted higher in the average representational consistency. A consistent representation or network discovery method will yield a low distance score, as networks that are exactly the same would receive a value of 0, whereas completely orthogonal networks would receive a value of 1.

To determine the optimal matching, we first calculate the cosine distance of all potential pairwise matches between supervised networks from each set of data. We then use the Hungarian Matching Algorithm ^64^ to determine the unique pairwise matching that minimizes the average cosine distance, meaning that each supervised network learned on the first set of data has a unique match to a supervised network learned on the second set of data. In other words, if network 1 from the first set of data is very similar to network 2 from the second set of data, those networks would be matched as they would have a low cosine distance.

#### Single-assay Model Formulation and Training

For single-assay model training, we isolated one of our three training assays (EPM, BOF, FLX) to use as our training dataset for dCSFA-NMF. While we focused on our model training using the FLX assay in the results section, we also trained models using the EPM and the BOF as singular training datasets. Each of our three single-assay models used the same labeling structure outlined above (see Results section entitled *"Distributed electome networks encode a convergent anxious internal state"*).

These models were learned using 4-fold cross validation on the training data. Models were selected based on predictive performance. Predictive performance was evaluated on the concatenated training and validation partitions of the other two assays not used for model training. It is worth noting that some bias exists in this evaluation, as some mice in the other two assays also experienced the third assay and their data may have been present in the training set of the selected model-training assay. However, even with this bias, these models failed to generalize across assays. The test, or holdout, partitions of all three assays were left untouched as each of the single-assay models failed to generalize to all three assays.

We performed our single-assay model training twice. First, we trained the models using a single supervised network^26^. Second, after we identified the value of multi-assay training and tuning for multiple consistent predictive networks, we reperformed our single-assay training with three supervised networks (as was selected in the multi-assay model below) and 27 unsupervised networks to allow for the most similar comparison with the multi-assay model. This second round of training was important as we wished to properly attribute whether generalization improvements from the multi-assay model came from increased model capacity or were due to the multi-assay training procedure.

#### Multi-Assay Model Formulation and Training

As mentioned above, our multi-assay model formulation involves concatenating the EPM, FLX, and BOF training datasets into a single training dataset. Since the labels for each assay are distilled into high- and low-anxiety states, data concatenation is possible because the labels are consistent. For multi-assay model training, we performed group 4-fold cross-validation on the training data partitions of all three assays for hyperparameter tuning. As mentioned earlier (*"Training and Test Splits"* section), all the data for each mouse is constrained to be completely with a training or validation fold. We then select model parameters based on the strategy described below in the *"Hyperparameter Selection Strategy"* section. The final model is learned on all training data with the selected hyperparameters, which is then used to evaluate on the test data.

#### Hyperparameter Selection Strategy

The dCSFA-NMF model requires selection of several hyperparameters. These factors include the number of electome networks *K*, number of supervised networks *Q*, the importance of the supervised task *λ*, and the importance of the supervised factor reconstruction *α*. These terms all impact the model learned as described in the "*Discriminative Cross-Spectral Factor Analysis (dCSFA-NMF)*" section. The number of electome factors *K* control how well we can reconstruct the original LFP data. The number of supervised networks *Q* impacts how well we can decode our anxiety states and our representational consistency. High representational consistency indicates that the same underlying network structure is reliably recovered across training perturbations, which is essential for interpreting the learned electome networks as stable, biologically meaningful circuits rather than dataset-specific solutions. In other words, across plausible data perturbations, the same circuit-level structure is repeatedly recovered. High representational consistency is required for downstream experimental design because causal perturbations necessarily target specific circuit features; without a stable network structure, experimental outcomes would be ambiguous, as null results could reflect instability of the learned representation rather than a true lack of biological relevance. Identifying a suitable number of supervised networks is especially crucial in the case where multiple underlying networks may be driving the emotional or behavioral state. Underspecifying the number of supervised networks to learn may result in the model inconsistently swapping across a subset of these suitable underlying networks.

We defined the encoder *f_θ_*(*X*) as a small neural network; specifically, it was a multi-layer perceptron with a single hidden layer of size 256 with a leaky rectified linear unit after the first hidden layer and a softplus nonlinearity before the output to maintain positivity of our network scores. We set *λ* = *α* = 1, and found our model performance was robust to changes in the network definition and those hyperparameters.

To choose the value of *K*, we first performed 4-fold cross-validation-over-subjects for *K* = {2, 4, ..., 58} while limiting the number of supervised networks *Q* = 1. Each model was trained on the training mice for all three assays jointly and evaluated on the validation mice for all assays, as per the multi-assay training procedure. We observed that the predictive performance of all three assays stabilized at *K* = 18 with little change across all three assays for higher values (Supplemental Fig. S5A). Subsequently, we found that the reconstruction performance plateaued at *K* = 30. Given that the predictive performance was consistent for *K* > 18, we selected *K* = 30 as the total number of networks that our model would learn. We evaluated this pattern for different values of *Q* and found the same patterns.

To choose a value of *Q*, we aimed to balance 3 design priorities in our model formation. First, our model must be predictive of the behavior of interest. Second, our model should find a relatively consistent solution (i.e., discovered brain networks should be similar across multiple runs), as this is necessary to have confidence in the networks and downstream experiments based on them. Lastly, our solution should be simple. Suppose we were to supervise all the networks in our model. We likely would achieve strong predictive performance; however, multiple-hypothesis testing problems would arise as we begin to test the relationships of each network with behavior. Therefore, we wished to find a stable, predictive solution that makes use of the smallest *Q* of supervised networks possible. To ascertain the value of *Q* we should use, we learned models with values *Q* = {1, 2, 3, 4, 5, 10, 20} with *K* = 30 using 4-fold cross-validation-over-subjects. We calculated the predictive performance on our anxiety assays and evaluated the average consistency score between the folds. The number of supervised networks does not greatly affect the overall prediction quality (Supplemental Fig. S5B, left), however, increasing the total number of supervised networks can significantly improve the representational consistency of the behaviorally relevant networks discovered (Supplemental Fig. S5B, right). We observed that the average AUC across all three assays peaks around *Q* ∈ {3, 4} and declined slightly as the number of supervised networks further increased, as shown in Supplemental Fig. S5A.

Additionally, we constrained our model to only identify supervised networks with scores that positively correlated with predicting a high-anxiety state, as we aimed to discover an anxiety network indicative of higher anxiety states rather than an anxiety inhibition network.

To learn the models, we first pretrained our factors by training a traditional nonnegative matrix factorization model (NMF) on our data. We selected stochastic gradient descent as our optimization algorithm as it is known to offer better generalization performance despite requiring longer training times ^65^. We used a learning rate of .001 and a momentum value of .9.

Our datasets were imbalanced with the longer-duration FLX neural recordings making up a much higher percentage of our total data. Therefore, we resampled windows from EPM and BOF such that each experimental context was equally influential on our network pretraining. We then froze the weights of our sorted NMF factors, trained the encoder to learn scores corresponding to the fixed factors, and trained the classifier to predict corresponding labels (high- and low-anxiety states) based on those scores. We found that training the encoder and classifier for 500 epochs of neural recording data (e.g., 500 passes over all training data) was sufficient for the optimization algorithm to converge and stabilize at a minimum of the loss function.

After pretraining, we then unfroze all parameters and trained them jointly. We found that an additional 500 epochs were sufficient for the model training to converge and stabilize at a minimum of the- loss function.

### Model Validation

#### Networks Decoded Assays Jointly

Given that our dCSFA-NMF model selection procedure identified three separate networks (see "*Hyperparameter Selection Strategy*"), we wished to validate that each of the learned networks are not simply learning to predict for each of the three training assays independently. Networks that truly capture anxiety should not be relevant to only one context where anxiety may be experienced, but rather should generalize to multiple contexts. Here, we evaluated whether the mean of the per-mouse AUC for each of the networks was different than the null value of 0.5 using a Mann-Whitney U test in each of the three training assays (Fig. 2D).

#### Individual Network Contribution Towards Prediction

We also considered whether one or more of our networks may not have contributed substantially to the overall prediction of the mouse’s internal anxiety state. To evaluate this, we considered the mean prediction logit of each network given by the mean network score multiplied by its corresponding logistic regression coefficient, and normalized it by the sum the mean prediction logits of all supervised networks. More formally, we defined the mean logit of an individual network as *z_j_* = *s̄_j_ϕ_j_*, where *s̄_j_* is the mean network activity score from the holdout data for network *j*, and *ϕ_j_* is the dCSFA logistic regression coefficient corresponding to that network activity score. Since we constrained our network to have positive network activity scores and logistic regression coefficients, no absolute value or squaring of the logits was necessary for comparison. We then evaluated the contribution of each network *j* as:

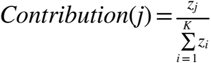

This metric quantified the average impact of each network towards predicting the anxiety context, as evaluated using holdout data.

#### Detailed Methods for Validation Analyses

##### Projecting Data into a Previously Learned Network

To evaluate network activity in datasets not used for model training (e.g., from sleep, delayed sucrose reward, optogenetic, fear conditioning, social preference, *ClockΔ19*, and stress experiments), we projected neural data into previously learned electome networks, i.e., *EN-Anxiety* and/or *EN-SocialAppetitive*. For each dataset, local field potential (LFP) features (power, coherence, and directionality) were extracted using the identical preprocessing and feature extraction pipeline described above. These features were then input into the trained dCSFA-NMF encoder, with all model parameters fixed, to compute network activity scores for each time window. No model retraining or parameter updating was performed during this step.

##### Location-based Dynamics in Network Activity

To compare post-exposure effects for the safe and avoidance zones in the EPM/BOF on *EN-Anxiety* dynamics, we extracted data from each timepoint in each location. We also extracted data in the five seconds preceding and following these timepoints (Figure 3). Here, the locations of C57 mice (closed arm/open arm/center for EPM and center/periphery for BOF) were encoded using Ethovision on 50 frames-per-second video recordings of the task, tracked at 25fps based on their center of mass. Frame labels were then aligned with our one-second resolution LFP features by assigning the label of each 1-s window that made up most of the Ethovision frames for that timepoint. For example, a time window was labeled as open arm if 20% of the Ethovision frames corresponded with the open arm location.

We then determined the *EN-Anxiety* activity at all instantaneous timepoints where the mouse was in the region of interest and at all observed timepoints within a +/- 5-second window from the instantaneous timepoint. Timepoints for which electrophysiological data, and therefore network scores, were not observed due to electrophysiological artifact, were dropped. This resulted in 10.5% of timepoints in the open arm case and 10.7% in the closed arm case being dropped. For each mouse, network activity was averaged within the -5 to -1s window, the 1s to 5s window, and the instantaneous timepoint window. Activity was then normalized to home cage activity levels and compared between locations across mice.

##### Fluoxetine Network Dynamics

We validated our network’s ability to reflect anxiety attenuation post-injection with saline or fluoxetine in the holdout mice from the FLX training task. Nine mice were injected with saline or fluoxetine at t=0 and we recorded neural activity for one hour post-injection. We then observed *EN-Anxiety* activity during the full hour recording for the 9 holdout mice. Network activity scores were binned and averaged at a 5-minute resolution with the mean activity (and standard error of the mean) across mice plotted in Fig. 3A. It is worth restating that our model was trained only using timepoints during the second half of the hour-long recorded data to ensure full absorption of the injected fluoxetine and to minimize handing effects. While our model was biased to distinguish between fluoxetine and saline due to our model training task, the model had no prior exposure to timepoints between t= [0,30] mins and no explicitly supervised trend for those timepoints. Data were analyzed using a two-way repeated measures ANOVA.

##### Delayed Sucrose Reward Task

We examined *EN-Anxiety* activity during the delayed sucrose reward assay to validate that *EN-Anxiety* was not encoding reward or arousal. Pump events for delivering sucrose to the mice were much shorter than our 1-second windows, therefore we used the event-triggered feature extraction code to precisely extract 1-second windows surrounding an event (sucrose delivery). We collected LFP features for 4 seconds prior to and 4 seconds after the pump event. These features were then projected into *EN-Anxiety*. Mean network activity at one second prior and one second post-pump event was then compared across mice (n=8) using a one-tailed Wilcoxon signed rank test (see *"Statistical Analysis Philosophy"* for rationale behind one-tailed test).

##### Optogenetic Stimulation of Ventral Hippocampus Circuits

We quantified network activity during optogenetic stimulation of the VHip → LH circuit and a control VHip → Amy circuit. We previously demonstrated that in the absence of ChR2, blue light stimulation has no direct impact on LFP activity using our recording approach^58^. Moreover, yellow light stimulation has no direct impact on LFP activity in the presence of ChR2^26,50^. Thus, we chose to compare network activity during yellow light stimulation and blue light stimulation, which all animals received. This approach enabled us to perform within-subject comparisons.

##### Fear Conditioning

For *EN-Anxiety* validation, event-centered features were extracted for 10 seconds prior to and 20 seconds after the tone onset. Mean *EN-Anxiety* activity was then calculated for the CS- and CS+ mice for the 10-second pre-segment and the 20-second post-segment separately. For both segments, we utilized a repeated measures two-way ANOVA to determine the impact of the 7^th^ and final tone/air puff event within and between groups. We isolated the 7^th^ event under the assumption that the CS+ mice had successfully associated the tone stimulus with the air puff across the 6 prior events. For analyzing the response of *EN-Anxiety* during the air puff alone, we only utilized mice for which network activity could be isolated for the complete 2 seconds of the air puff (7/13 CS+ mice and 15/17 CS-mice).

### Statistical Analysis Philosophy

We trained a multi-region, multi-assay model to putatively encode the anxiety state that converged across multiple assays. We also trained models to test whether this putative state was encoded by single regions and/or by pairs of brain regions. We assessed each single- or paired-region model independently for the three assays. Based on our prior observations that other emotional internal states could not be decoded from individual brain regions ^26,27,51^, we hypothesized that activity from single regions and/or pairs of regions would fail to decode a convergent anxiety state as well. To increase the likelihood of falsifying our hypothesis, we chose to leave all statistical analysis using single region/pairs of regions uncorrected. We observed that P>0.05 for at least one assay for each single region/pairs of regions test (Fig. 1F, Supplemental Figure S6). Since correcting for multiple comparisons would have served to further increase the P-values, we concluded that no single region/pairs of regions encoded a convergent anxiety state.

Next, to validate the multi-region, multi-assay networks, we recorded LFP data in the same brain regions from new mice and/or new paradigms. We focused this subsequent analysis on Electome Network 2/*EN-Anxiety* since it independently encoded all three of our initial anxiety assays, it showed the highest contribution to the joint predictive model, and it did not represent a superposition of the FLX and BOF/EPM assays. Validation of *EN-Anxiety* involved various statistical tests and procedures. For comparison of high- vs. low-anxiety states, we performed non-parametric statistical tests on mean network scores for time segments or groups of interest. In cases where parametric tests are useful, such as examining network dynamics over time, we performed a Box–Cox transformation of the network activity scores prior to statistical testing. For the validation tests, we expected the activity of *EN-Anxiety* to be higher in the high-anxiety state compared to the low-anxiety state since the learned model matched higher network activity to the high-anxiety contexts. For these cases, we performed statistical tests with a one-tailed test. A similar approach was implemented for our control network analyses. Specifically, we tested whether network activity responded in the same manner to other psychological constructs like arousal and reward as it had to anxiety.

We hypothesized that anxiety-related behavior and network activity would be lower in the *ClockΔ19* mouse model of mania. Thus, we utilized a one-tailed test for behavior. However, we utilized a two-way ANOVA for network analyses since we were testing the impact of both genotype and exposure to the EPM assay on network activity, and we had no *a priori* hypotheses about these effects. All such cases are disclosed in our results section. All p-values are reported as uncorrected p-values for network analyses.

In some cases, each mouse has data windows missing at different nonrandom time points. Indeed, as observed when we examined network dynamics in the safe regions over time (Fig. 3B), each mouse leaves the safe regions at different time points throughout the 5-minute session. Therefore, for this analysis, we make use of an ANCOVA-based strategy, since it is flexible with missing data and allows analysis of dynamics over time. A disadvantage of this approach is that samples are treated independently without accounting for subject identity.

### Visualization

Networks were visualized as chord plots using code adapted from <https://github.com/carlson-lab/lpne/> to allow for recoloring of frequency bands. We visualize networks by measuring their relative expression compared to all other networks. Mathematically, this is expressed as the percentage of the reconstructed neural activity that can be attributed to each network. This relative expression was evaluated on the training mice; given the unsupervised nature of the reconstruction task, there is no meaningful difference between the training and testing mice. This strategy results in an even appraisal of low- and high-frequency features, even though low-frequency spectral features tend to have higher power. We then selected the 85^th^ percentile of these contributions, which is a threshold that adequately highlights dominant features without cluttering the plot and is consistent with related works ^26^. Finally, these features were plotted with a transparency ranging from 40 to 90% based on their strength with each network.

### Reproducibility

#### Computational Environment and Codebase Disclosure

Preprocessing and feature extraction code was performed in MATLAB R2022a using the LFP feature extraction pipeline found on the main branch at https://github.com/carlson-lab/lpne-data-analysis. Event-triggered feature extraction code can be found on the "framewindows" branch of the same repository. A PyTorch implementation of dCSFA-NMF can be found at https://github.com/carlson-lab/lpne/. All code for network generation, hyperparameter tuning, model implementation, plotting, and a singularity definition file for replicating our Python environment can be found https://github.com/carlson-lab/Anxiety. Development was performed on a computer cluster in a Singularity Container managed Python environment with nodes utilizing an NVIDIA RTX 2080 Ti.

## ACKNOWLEDGEMENTS

We would like to thank Elise Adamson, Jake Benton, and Rachel Fisher-Foye for technical assistance, and Karim Abdelaal for comments. A special thanks to Freeman Hrabowski, Robert and Jane Meyerhoff, and the Meyerhoff Scholarship Program.

## AUTHOR CONTRIBUTIONS

Conceptualization DNH, MHK, RCH, SDM, DEC and KD; Methodology, DNH, MHK, WEC, DEC, and KD; Formal Analysis, DNH, MHK, KKWC, GET, YG, YF, MT, and KD; Investigation, DNH, MHK, KKWC, GET, YG, DW, AM, YF, MT, AF, NMG, and KD; Resources, CAM, RCH, DEC and KD; Writing – Original Draft, DNH, MHK, JMZ, DEC, and KD; Writing – Review & Editing, DNH, MHK, KKWC, CAM, JMZ, RH, SDM, DEC, and KD; Visualization, DNH, MHK, DEC, and KD; Supervision, CAM, RCH, DEC and KD; Project Administration, DEC and KD; and Funding Acquisition, DNH, RCH, DEC, and KD; see Supplemental Table S1 for experiment-specific contributions.

## DATA AND CODE AVAILABILITY

Code for LFP feature extraction, model training, and network visualization is available at the following repositories: the LFP feature-extraction pipeline (MATLAB R2022a) at https://github.com/carlson-lab/lpne-data-analysis (main branch; event-triggered feature extraction on the “framewindows” branch); the PyTorch implementation of dCSFA-NMF at https://github.com/carlson-lab/lpne/; and code for network generation, hyperparameter tuning, model implementation, plotting, and a Singularity definition file for replicating the Python environment at https://github.com/carlson-lab/Anxiety. [Author: please add a repository name/accession number for the raw electrophysiological and behavioral data, if one is available, e.g. Dryad, Zenodo, or Figshare – none was stated in the manuscript text provided.]

