## Supplemental Materials for "An electrical brain network encodes anxiety in health and depressive states"

*An Electome Network Encodes Anxiety*

### SUPPLEMENTARY TABLE S1: DETAILED AUTHOR CONTRIBUTIONS

| Author | Contribution |
|---|---|
| **Dalton N. Hughes** | Designed and performed FLX, EPM, BOF, VHip → LH ChR2, *ClockΔ19*, and chronic mild unpredictable stress EPM experiments. Oversaw data collection for fear conditioning experiment. Conceived the multi-assay machine learning approach with MHK. Secured funding and resources. Prepared original figures. Performed statistical analysis. Wrote original draft of the introduction, results, materials and methods and discussion. Revised manuscript. |
| **Michael Hunter Klein** | Performed feature extraction and preprocessing. Performed single-assay modeling of FLX, EPM, and BOF datasets. Conceived and performed the multi-assay machine learning approach with DNH. Performed all machine learning analyses and developed codebase utility for multi-assay learning. Developed multiple network and stability validation approach in consult with WEC and DEC. Performed all dataset projections and visualizations of learned electome networks. Verified all statistical analysis. Wrote original draft of the introduction, results, materials and methods and discussion. Revised manuscript. |
| **Kathryn Katsue Walder-Christensen** | Implanted mice for initial FLX, EPM, and BOF experiments in male mice, EPM experiments in female mice, *ClockΔ19* mice experiment, and fear conditioning experiment. Performed chronic mild unpredictable stress behavioral induction paradigm. Performed neurophysiological recordings for the chronic mild unpredictable stress experiment. Performed video tracking and behavioral analysis for EPM experiments in implanted C57 and *ClockΔ19* mice. Contributed to histological analysis for mice used in initial FLX, EPM, and BOF experiments, chronic unpredictable mild stress experiment, and fear conditioning experiment. Revised manuscript. |
| **Gwenaëlle E. Thomas** | Performed viral injections and histology validation for VHip → LH ChR2 experiment. Designed FLX experiments with DNH. |
| **Yael Grossman** | Performed chronic social defeat stress paradigm and gathered data for neurophysiological analysis with RCH. Implanted animals for fear conditioning experiment. Contributed to histological analysis for chronic unpredictable mild stress experiment and fear conditioning experiment. |

| | |
|---|---|
| **Diana Waters** | Performed EPM and BOF neurophysiological experiments. |
| **Anna Matthews** | Performed EPM behavioral experiment in fluoxetine-treated mice. |
| **William E. Carson** | Developed a new reconstruction loss formulation for supervised networks. Consulted on theory regarding network stability procedure with DEC. |
| **Yassine Filali** | Performed EPM behavioral experiment for chronic social defeat stress paradigm. |
| **Mariya Tsyglakova** | Performed behavioral testing in unimplanted *ClockΔ19* mice and their littermate controls. |
| **Alexandra Fink** | Coordinated and performed chronic unpredictable mild stress behavioral paradigm. |
| **Neil M. Gallagher** | Coordinated and performed chronic unpredictable mild stress behavioral paradigm. |
| **Masiel Perez-Balaguer** | Performed background review, contributed to initial manuscript draft. |
| **Colleen A. McClung** | Provided *ClockΔ19* mouse line, oversaw behavioral testing in unimplanted mice, and revised the paper. |
| **Jean Mary Zarate** | Designed approach to integrate neurophysiological and behavioral analyses with KD, wrote and edited the manuscript. |
| **Rainbo C. Hultman** | Designed and performed VHip → LH ChR2 experiment with SDM and GET. Performed chronic social defeat stress paradigm and gathered data for neurophysiological analysis with YG. Oversaw collection of chronic social defeat stress behavioral data. |
| **Stephen D. Mague** | Implanted mice for network discovery and validation using FLX, EPM, and BOF. Implanted *ClockΔ19* mice and littermate controls. Designed and performed VHip → LH ChR2 experiment with DNH and RCH. Performed neural recordings in female C57 mice. Revised the manuscript. |
| **David E. Carlson** | Supervised and oversaw all experiments and analyses with KD. Secured resources, wrote and revised the manuscript. |
| **Kafui Dzirasa** | Supervised and oversaw all experiments and analyses with DEC. Implanted mice for VHip → LH ChR2 experiment. Secured resources, wrote and revised the manuscript. |

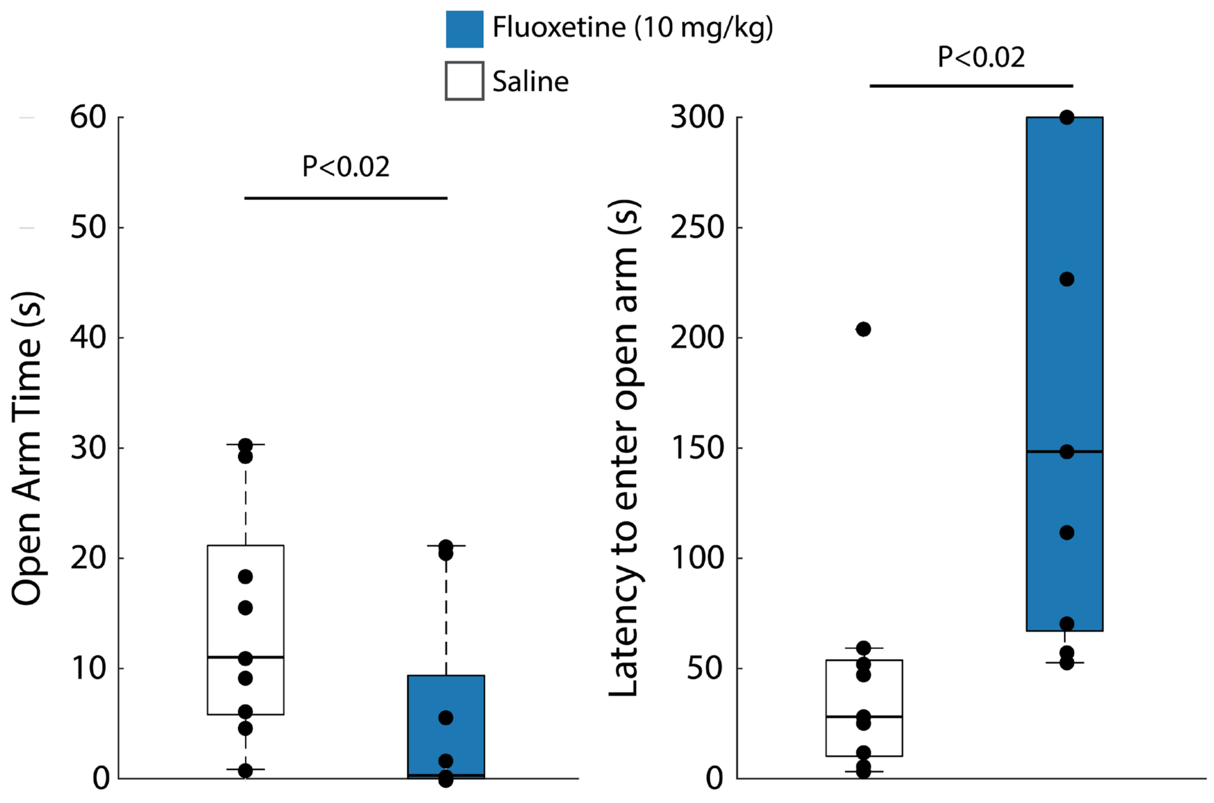

**Supplemental Figure S1: Behavioral impact of acute fluoxetine treatment in unimplanted male mice.** Mice were treated with 10mg/kg fluoxetine or saline in their home cage, and behavior was assayed on an elevated plus maze for five minutes. Behavioral testing occurred 30 minutes after drug treatment (U=62, P<0.02; U=52, P<0.02, for open arm time and latency to entry, respectively; one-tailed Wilcoxon Rank-Sum test; N=9 mice/group). For the box and whisker plots, the central mark is the median, the edges of the box are the 25th and 75th percentiles, the whiskers extend to the most extreme datapoints the algorithm does not consider to be outliers, and the outliers are plotted individually as "+."

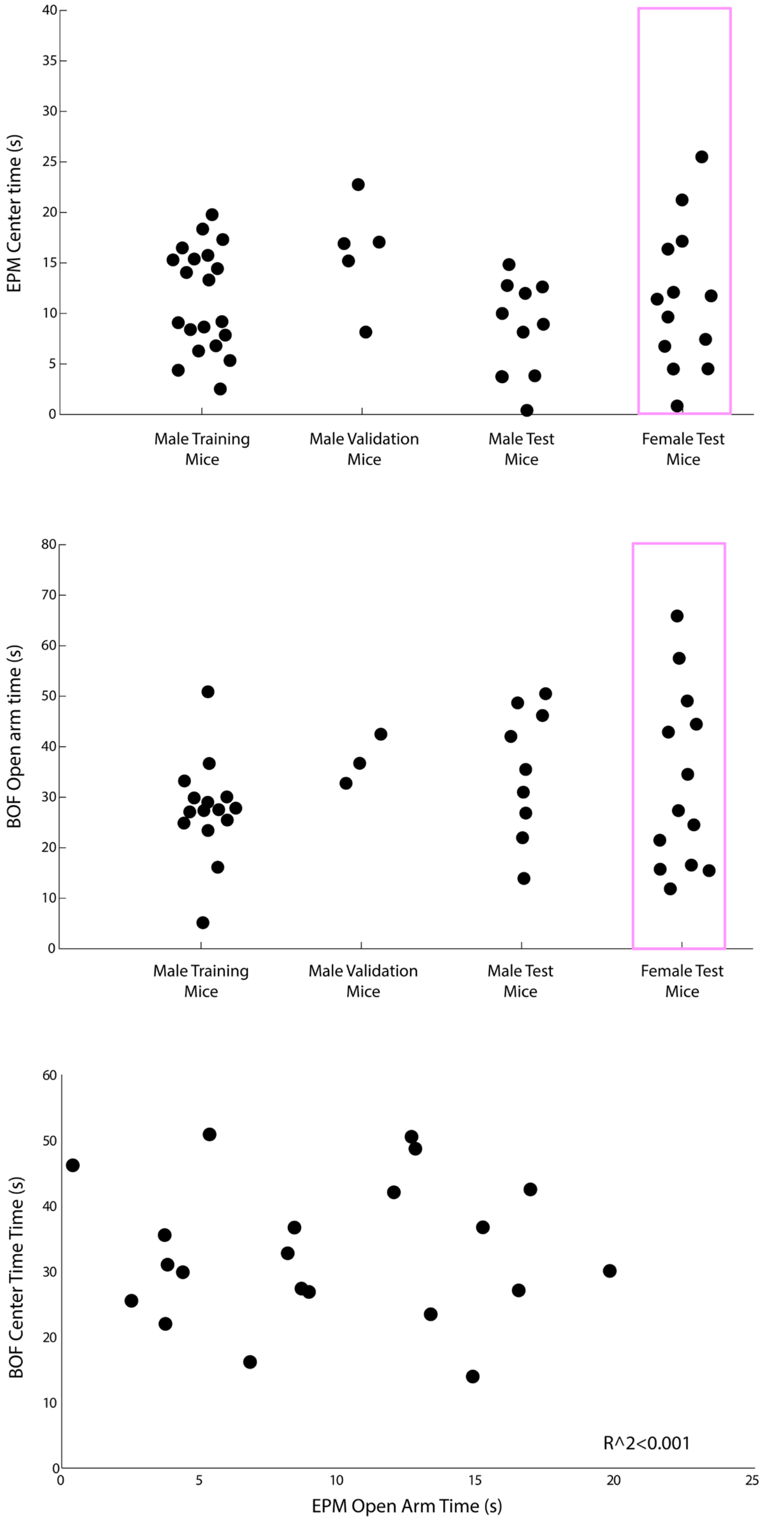

**Supplemental Figure S2: Avoidance behavior in male/female mice used to train, validate, and test anxiety network models.** EPM (top; N=21, 5, 11, and 13, respectively) and BOF (middle, N=15, 3, 9, and 13 respectively) avoidance behavior is shown for the training, validation, and test mice. Female mice are highlighted by the pink box. The correlation between these measures in male mice subjected to both assays (N=21 total training and test mice) is shown on the bottom ($R^2<0.001$ using Pearson correlation).

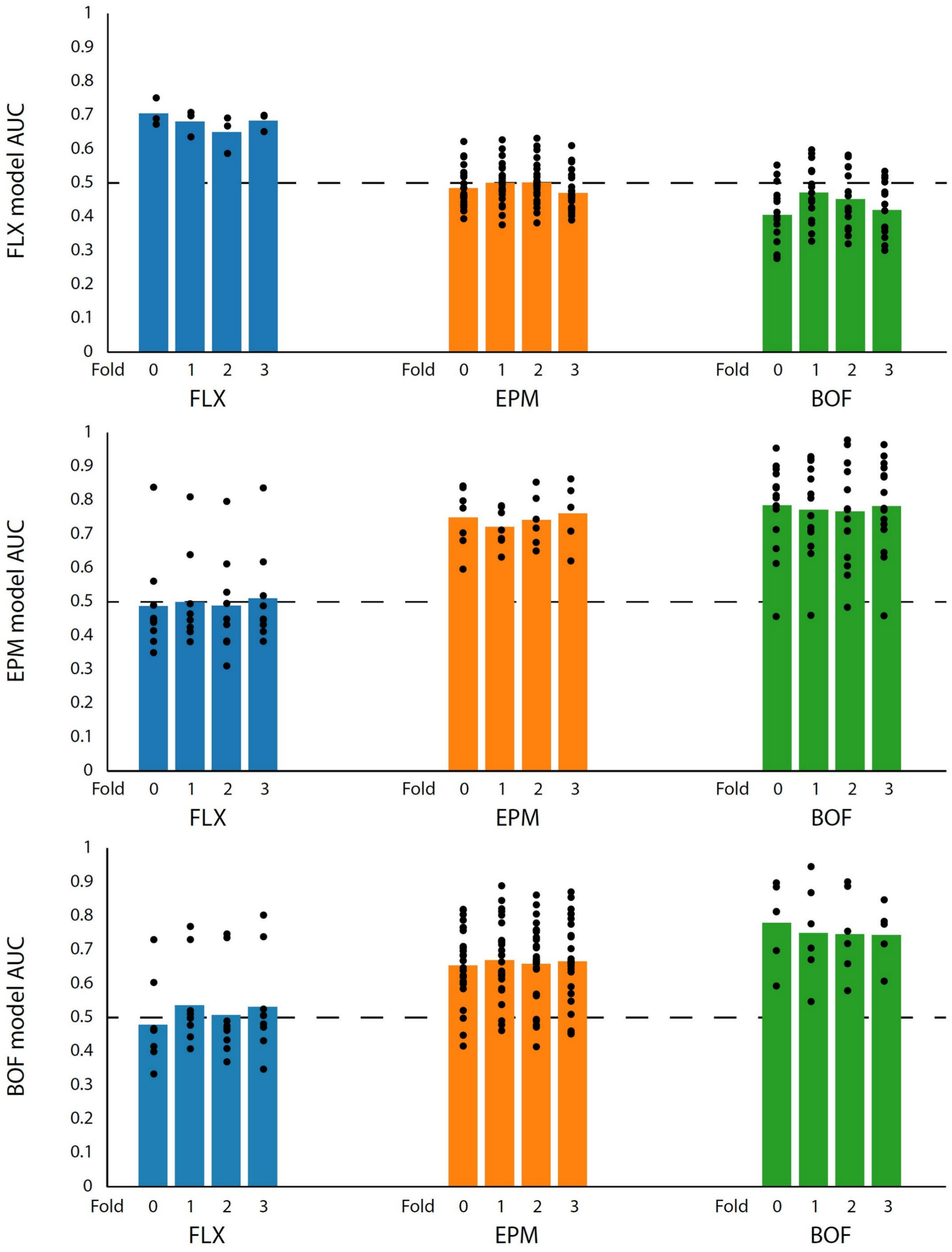

**Supplemental Figure S3:** Four-fold cross-validation results for models trained on each of the three assays separately (3-7 holdout mice within task per fold, and 9-26 holdout mice between tasks per fold). K-fold data splits were done using only the training and validation data of the final model. The number of data points for predictive performance on the training assay are reduced due to train-test splits. Dashed line at AUC = 0.5 corresponds to models with no predictive utility. These models solely utilized data from the initial 41 implanted male mice.

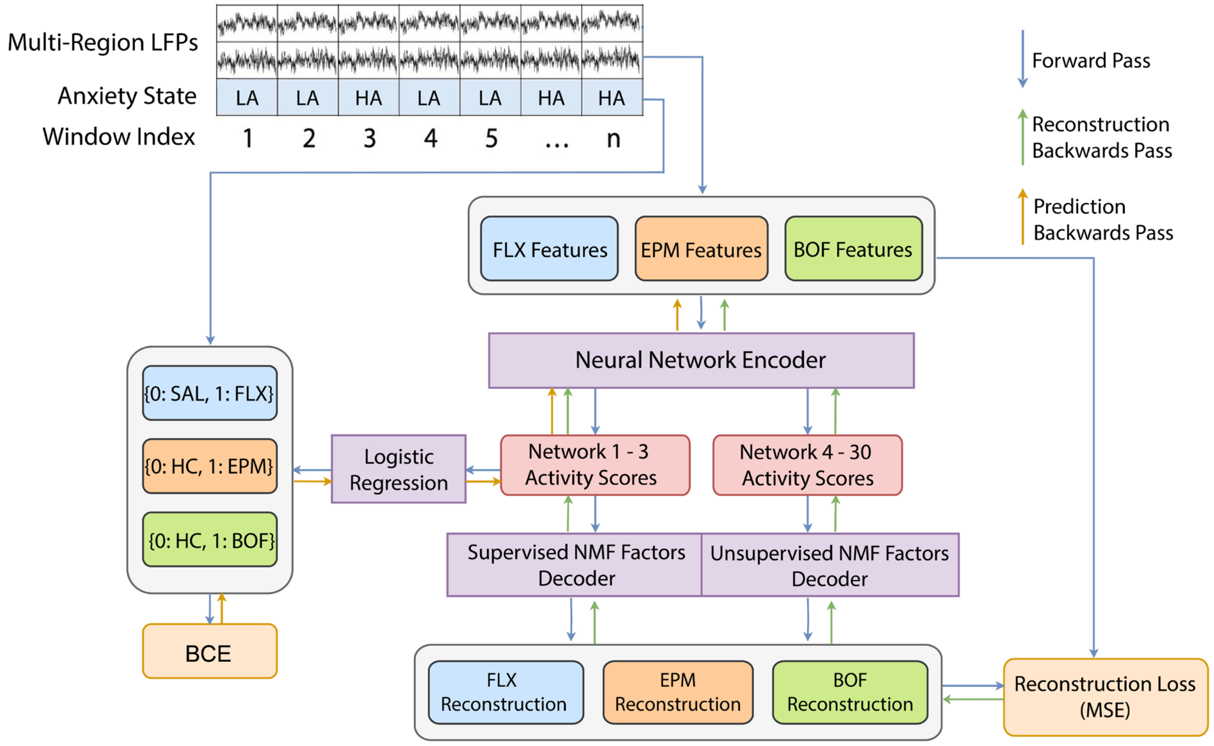

**Supplemental Figure S4: Detailed multi-assay network architecture for learning and validating anxiety-related electome networks.** Local field potential (LFP) recordings from multiple brain regions are segmented into fixed-length time windows and labeled as low-anxiety (LA) or high-anxiety (HA) states according to context (top). For each window, assay-specific feature sets are extracted from the multi-region LFPs recorded during the fluoxetine (FLX), elevated plus maze (EPM), and bright open field (BOF) paradigms. These features are passed through a shared neural network encoder that maps them into latent electome network activity scores (middle). A subset of networks is supervised to predict the anxiety state via logistic regression with a binary cross-entropy loss (BCE; middle left), while additional unsupervised networks capture shared structure across assays. In parallel, decoder pathways reconstruct assay-specific features from network activity (bottom), and reconstruction error (mean squared error, MSE) is used to regularize learning. Forward passes support anxiety state prediction, while backward passes provide gradients that are used to jointly optimize classification and reconstruction objectives. Note that backward passes from the supervised objective only directly impact activity score learning of the supervised networks (in this study, one to three networks) and the corresponding part of the neural network encoder.

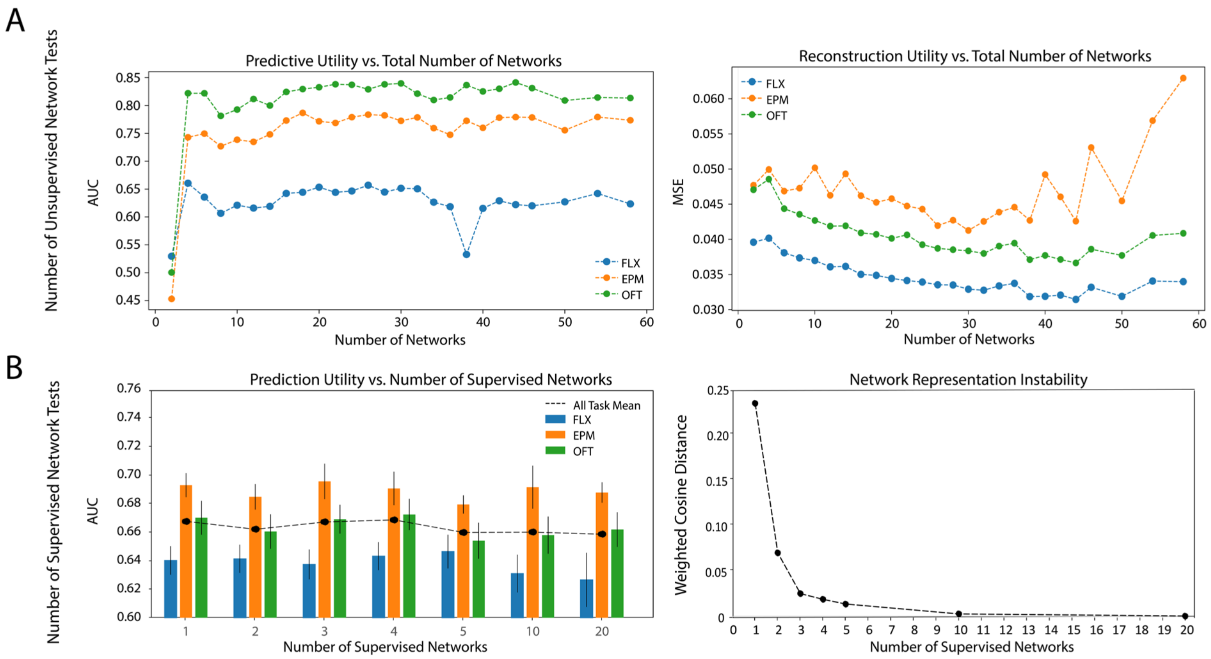

**Supplemental Figure S5: Hyperparameters for model selection. A)** Plots show predictive (left) and reconstruction performance (right) from total number of networks using grid search cross-validation for the multi-assay model. The number of supervised networks was fixed at one. We set the best number of total (supervised and unsupervised) electome networks to 30 to balance reconstruction and predictive performance. **B)** Plots show prediction performance and network representation instability across varying numbers of supervised networks using grid search cross-validation. Data shown as mean±s.e.m (left). N=3 supervised networks was selected as the optimal balance between prediction and representational stability using the elbow method (right, see Methods).

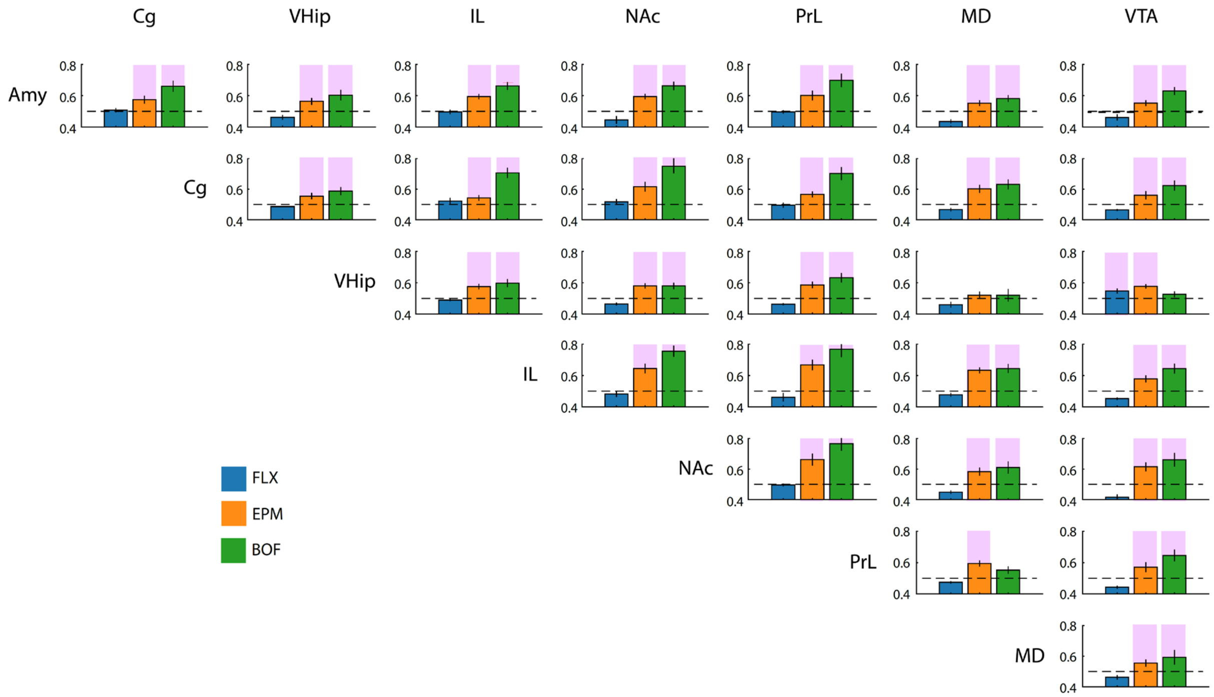

**Supplemental Figure S6: Individual neural circuits fail to independently encode a generalized anxiety state.** We trained dCSFA-NMF models using LFP features from pairs of brain regions (power, coherence and directionality) and tested them across the three anxiety paradigms used for training. The pairs of regions that contributed to each model are shown to the left and to the top of each plot. Tests were performed using 17 holdout mice, and pairs of regions that showed significant encoding are highlighted in pink (one-tailed unpaired t-test against a null AUC distribution at α = 0.05; dashed line corresponds to an AUC = 0.5). Data shown as mean±s.e.m. Note that while many pairs encoded two of the anxiety paradigms, none of them generalized to all three.

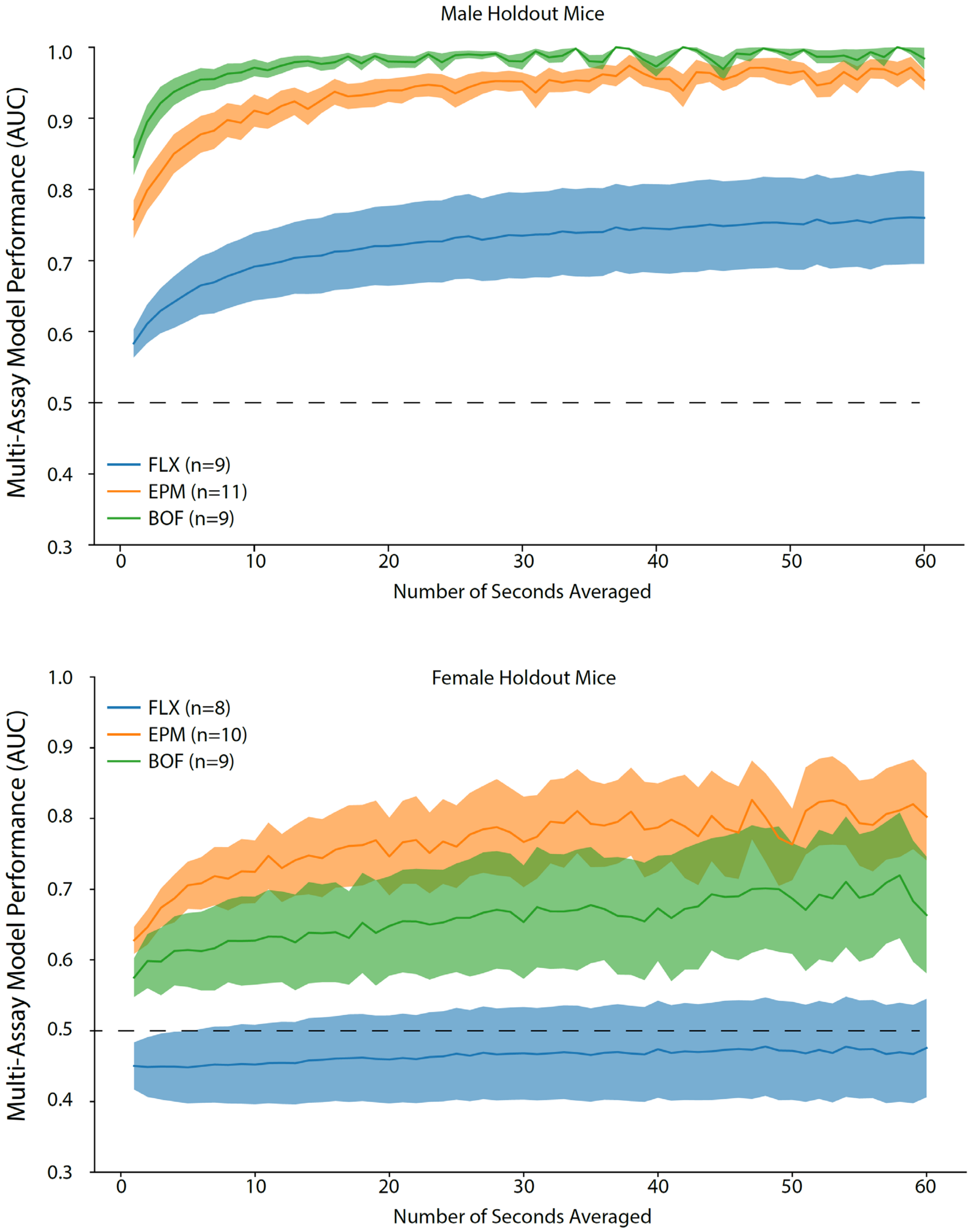

**Supplemental Figure S7: Multi-assay network model generalization to new male and female mice across larger data windows.** The dotted line at AUC=0.5 represents no information encoding. Data is shown as mean±s.e.m.

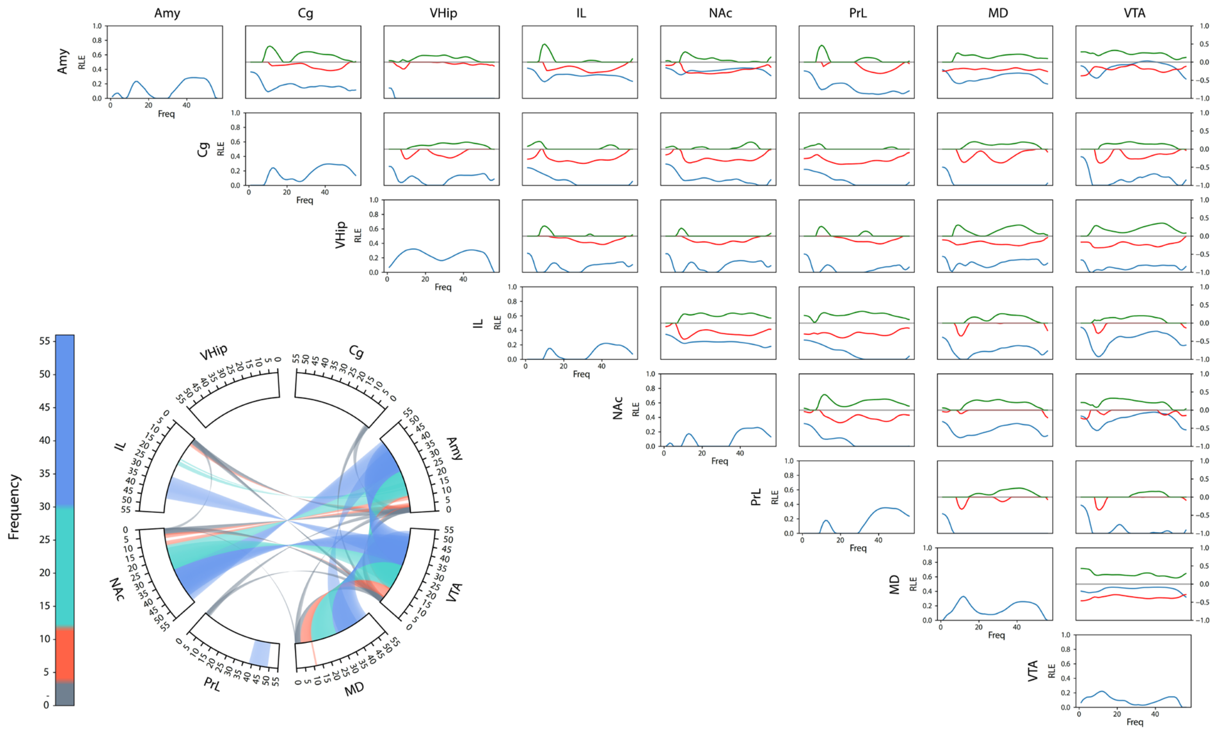

**Supplemental Figure S8: Anxiety Electome Network 1: Power, coherence, and directionality measures that define *Electome Network 1.*** Brain areas are shown to the top and the left of the rectangular graphs, identifying pairs of regions for which power and coherence, and directionality density functions are shown for Electome Network 1. Amplitude values (blue lines) reflect the relative LFP spectral energy (RLE) observed at each frequency, where the electome network is normalized to the total energy observed across all the learned networks in the multi-task model. The directionality functions for each pair of regions (A → B and B → A) are shown in red and green lines, respectively (axis scale to the right). Positive spectral offsets in green correspond to the directionality from the area listed on the top to the area listed on the left. Negative spectral offsets in red correspond to the directionality from the area listed on the left to the area listed on the top. The circular plot depicts the frequencies for power (outer rim) and coherence (curved lines connecting two regions) above an amplitude threshold of 0.33, corresponding to the top 15% of features.

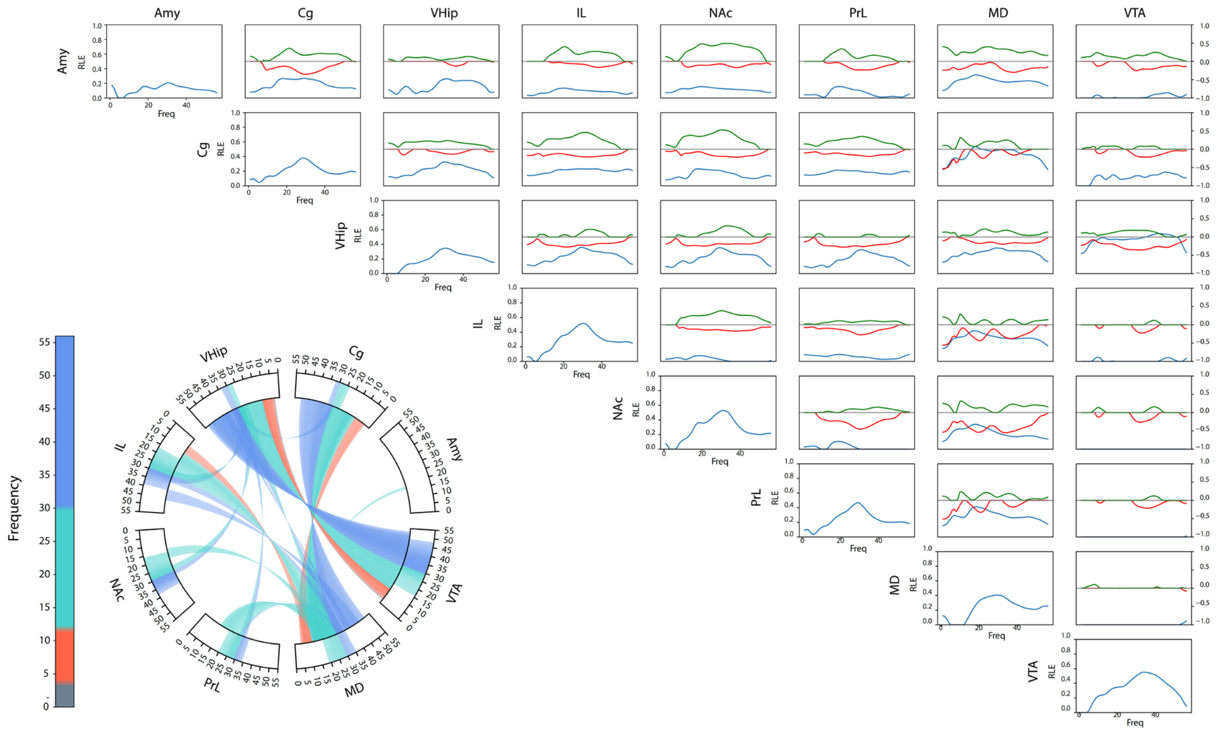

**Supplemental Figure S9: Anxiety Electome Network 2: Power, coherence, and directionality measures that define *Electome Network 2 (EN-Anxiety)*.** Brain areas are shown to the top and the left of the rectangular graphs, identifying pairs of regions for which power, coherence and directionality density functions are shown for Electome Network 2. Amplitude values (blue lines) reflect the relative LFP spectral energy (RLE) observed at each frequency, where the electome network is normalized to the total energy observed across all the learned networks in the multi-task model. The directionality functions for each pair of regions (A → B and B → A) are shown in red and green lines, respectively (axis scale to the right). Positive spectral offsets in green correspond to the directionality from the area listed on the top to the area listed on the left. Negative spectral offsets in red correspond to the directionality from the area listed on the left to the area listed on the top. The circular plot depicts the frequencies for power (outer rim) and coherence (curved lines connecting two regions) above an amplitude threshold of 0.31, corresponding to the top 15% of features.

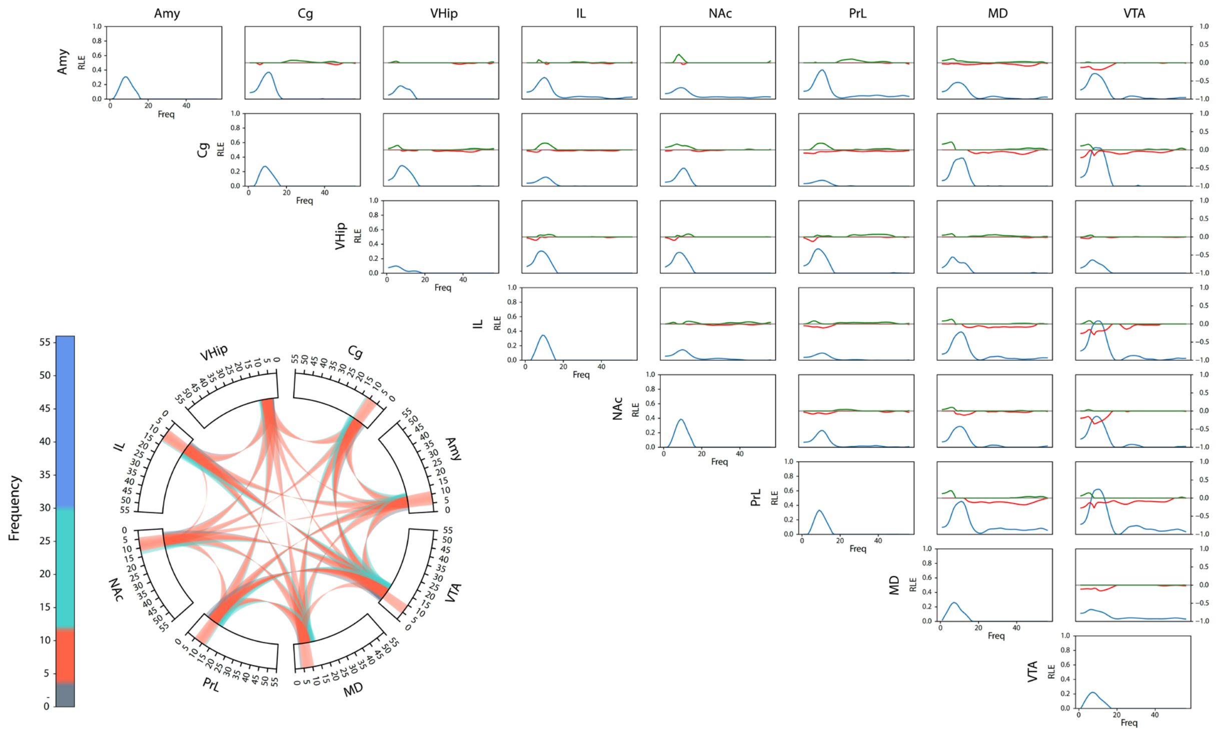

**Supplemental Figure S10: Anxiety Electome Network 3: Power, coherence, and directionality measures that define *Electome Network 3.*** Brain areas are shown to the top and the left of the rectangular graphs, identifying pairs of regions for which power, coherence, and directionality density functions are shown for Electome Network 3. Amplitude values (blue lines) reflect the relative LFP spectral energy (RLE) observed at each frequency, where the electome network is normalized to the total energy observed across all the learned networks in the multi-task model. The directionality functions for each pair of regions (A → B and B → A) are shown in red and green lines, respectively (axis scale to the right). Positive spectral offsets in green correspond to the directionality from the area listed on the top to the area listed on the left. Negative spectral offsets in red correspond to the directionality from the area listed on the left to the area listed on the top. The circular plot depicts the frequencies for power (outer rim) and coherence (curved lines connecting two regions) above an amplitude threshold of 0.13, corresponding to the top 15% of features.

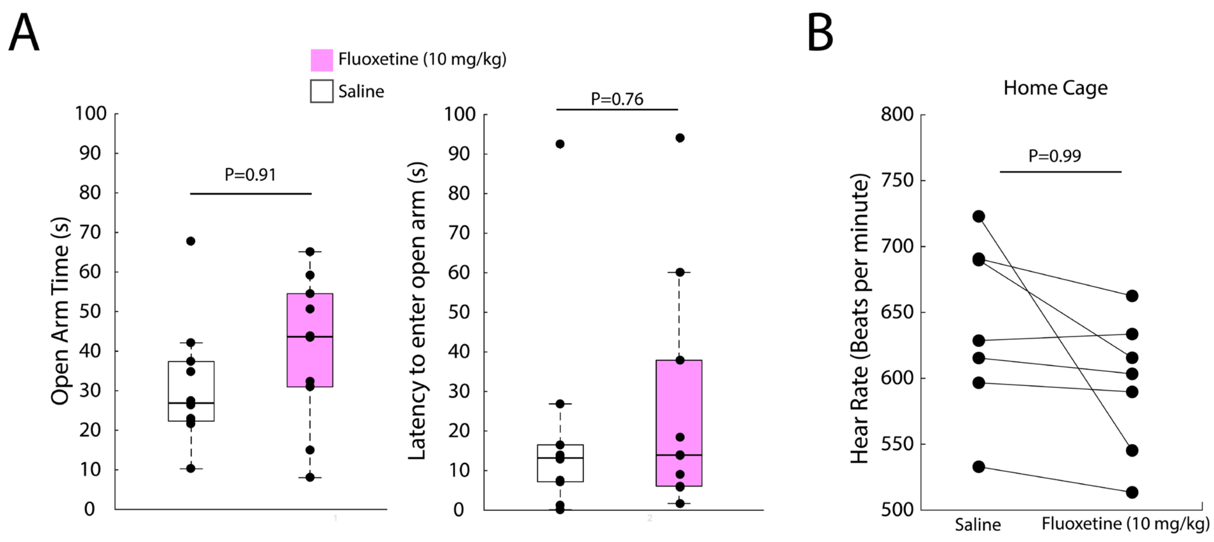

**Supplemental Figure S11: Behavioral impact of acute fluoxetine treatment in unimplanted female mice.** Mice were treated with fluoxetine 10mg/kg or saline in their home cage. **A)** Behavior was assayed on an elevated plus maze for five minutes. Behavioral testing occurred 30 minutes after drug treatment (N=9 mice/group). Data was analyzed using a one-tailed rank-sum test (U=122, P=0.91; U=114, P=0.76, for open arm time and latency to entry, respectively, using one-tailed Wilcoxon rank sum test). **B)** Heart rate was monitored for 60 minutes after saline and fluoxetine treatment in another group of mice (N=7 mice). The second 30 minute interval was utilized for physiological analysis, paralleling the neural recordings (U=27, P=0.99 using one-tailed sign rank test). For the box and whisker plots, the central mark is the median, the edges of the box are the 25th and 75th percentiles, the whiskers extend to the most extreme datapoints the algorithm does not consider to be outliers, and the outliers are plotted individually as "+."

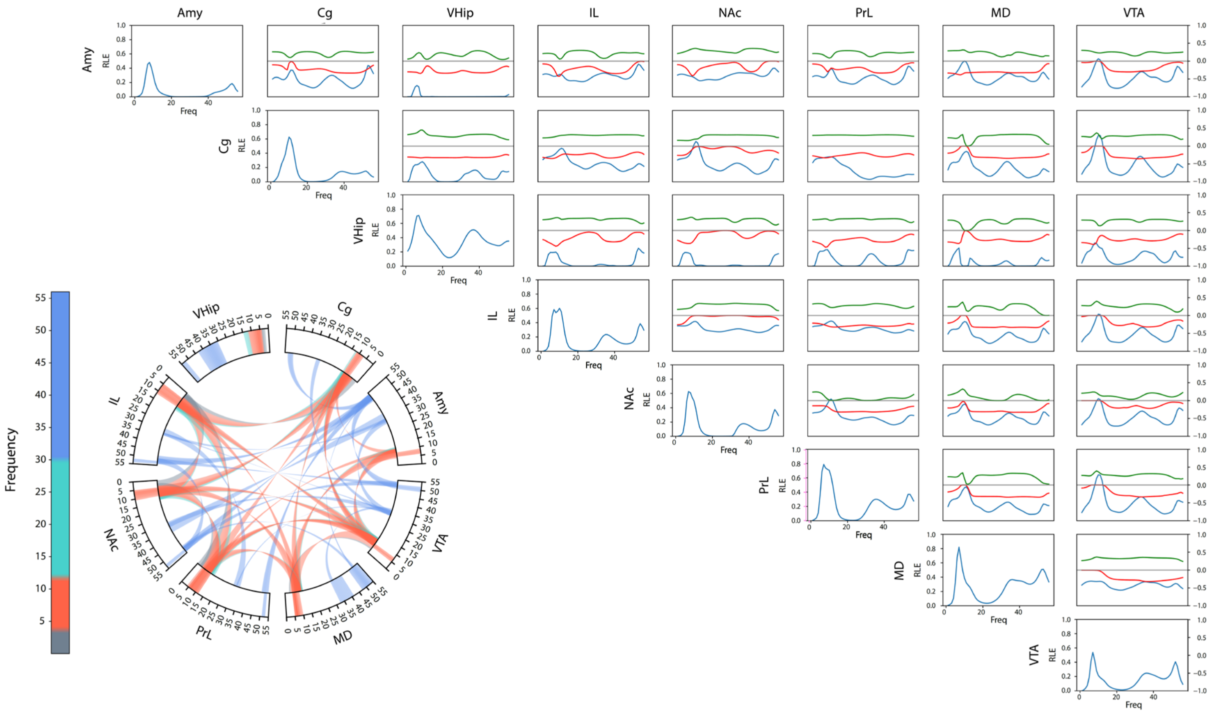

**Supplemental Figure S12: Social Appetitive Electome Network 3: Power, coherence, and directionality measures that define *Electome Network SocialAppetitive* (*EN-SocialAppetitive*).** This network was discovered and described in our prior work[1]; figure adapted from Mague et al.[1]. Brain areas are shown to the top and the left of the rectangular graphs, identifying pairs of regions for which power, coherence, and directionality density functions are shown for *EN-SocialAppetitive*. Amplitude values (blue lines) reflect the relative LFP spectral energy (RLE) observed at each frequency, where the electome network is normalized to the total energy observed across all the learned networks in the multi-task model. The directionality functions for each pair of regions (A → B and B → A) are shown in red and green lines, respectively (axis scale to the right). Positive spectral offsets in green correspond to the directionality from the area listed on the top to the area listed on the left. Negative spectral offsets in red correspond to the directionality from the area listed on the left to the area listed on the top. The circular plot depicts the frequencies for power (outer rim) and coherence (curved lines connecting two regions) above an amplitude threshold of 0.33, corresponding to the top 15% of features.

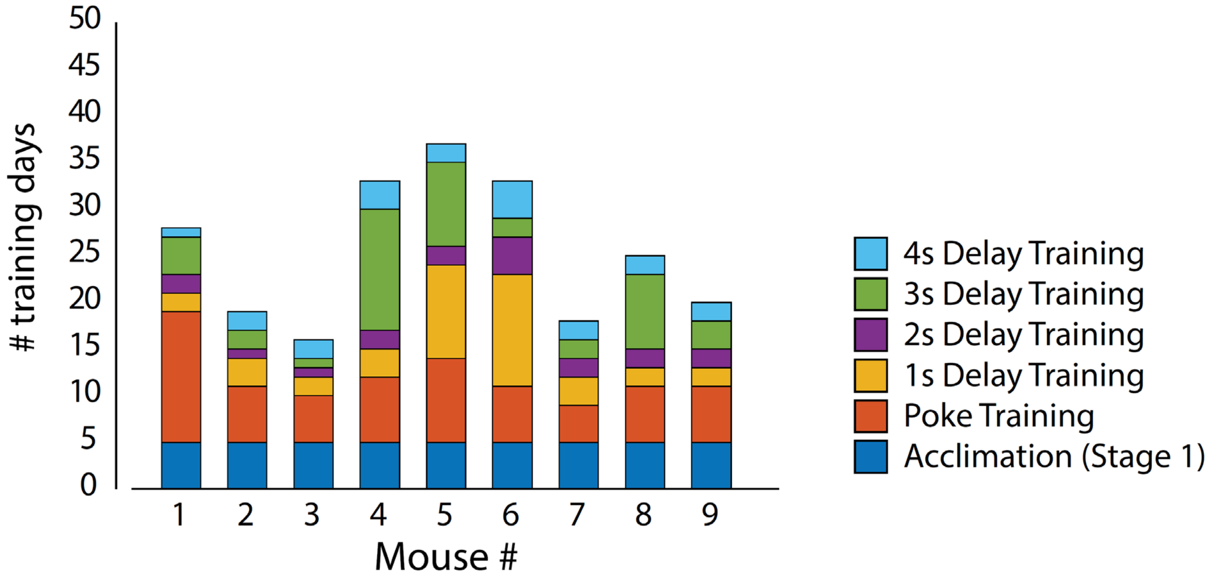

**Supplemental Figure S13: Number of days mice required to complete stages of training for reward task.** See Methods for details about the training protocol ("Delayed sucrose reward apparatus, training, and task").

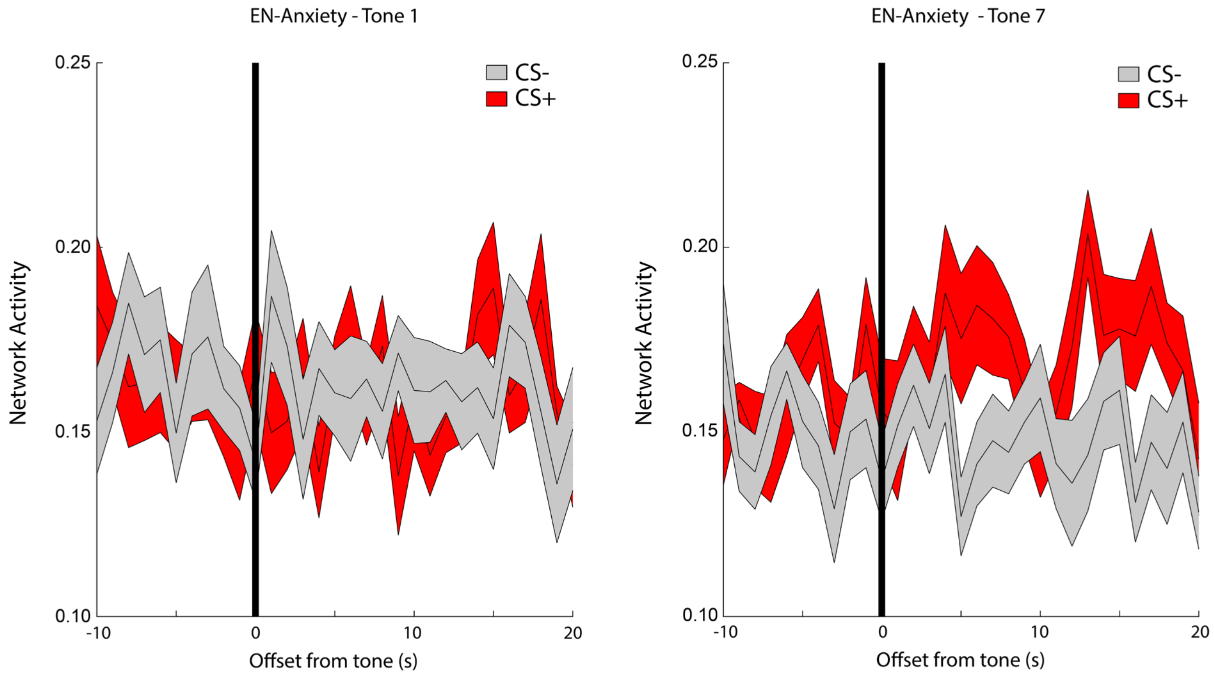

**Supplemental Figure S14: *EN-Anxiety* activity during fear conditioning paradigm.** *EN-Anxiety* activity was quantified relative to tone presentation. The vertical black line shows the tone onset. Plots show activity relative to the first (left) and last tone (right, same as Fig. 5M) in conditioned (CS+, red; N=13 mice) and non-conditioned control mice (CS-, grey; N=17 mice). Data shown as mean±s.e.m.

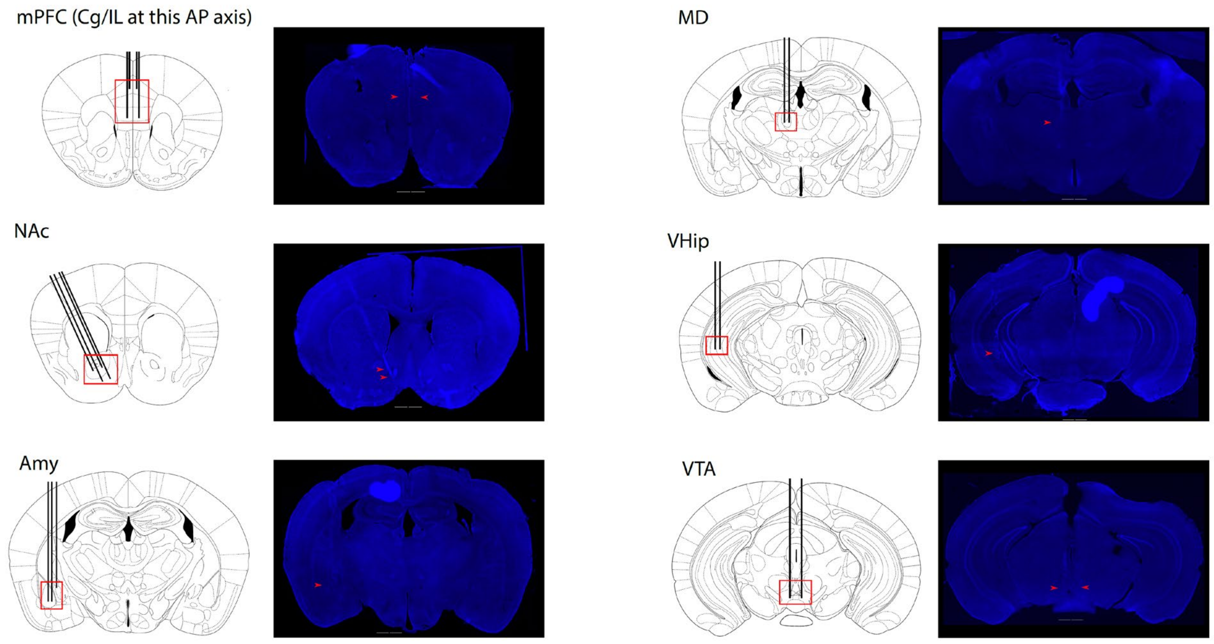

**Supplemental Figure S15: Histological confirmation of electrode placements.** Representative slices shown from the targeted brain regions, which include medial prefrontal cortex (mPFC; cingulate -Cg, prelimbic -PL, and infralimbic -IL cortex), nucleus accumbens (NAc), amygdala (Amy), medial dorsal thalamus (MD), ventral hippocampus (VHip), and ventral tegmental area (VTA). Red squares delineate the boundaries of electrode locations within each region, and red arrows highlight tips of implanted microwires in the representative brain slice.
